# A strategy for mapping biophysical to abstract neuronal network models applied to primary visual cortex

**DOI:** 10.1101/2021.04.28.441749

**Authors:** Anton V. Chizhov, Lyle J. Graham

## Abstract

A fundamental challenge for the theoretical study of neuronal networks is to make the link between complex biophysical models based directly on experimental data, to progressively simpler mathematical models that allow the derivation of general operating principles. We present a strategy that successively maps a relatively detailed biophysical population model, comprising conductance-based Hodgkin-Huxley type neuron models with connectivity rules derived from anatomical data, to various representations with fewer parameters, finishing with a firing rate network model that permits analysis. We apply this methodology to primary visual cortex of higher mammals, focusing on the functional property of stimulus orientation selectivity of receptive fields of individual neurons. The mapping produces compact expressions for the parameters of the abstract model that clearly identify the impact of specific electrophysiological and anatomical parameters on the analytical results, in particular as manifested by specific functional signatures of visual cortex, including input-output sharpening, conductance invariance, virtual rotation and the tilt after effect. Importantly, qualitative differences between model behaviours point out consequences of various simplifications. The strategy may be applied to other neuronal systems with appropriate modifications.

**Author summary:** A hierarchy of theoretical approaches to study a neuronal network depends on a tradeoff between biological fidelity and mathematical tractibility. Biophysically-detailed models consider cellular mechanisms and anatomically defined synaptic circuits, but are often too complex to reveal insights into fundamental principles. In contrast, increasingly abstract reduced models facilitate analytical insights. To better ground the latter to the underlying biology, we describe a systematic procedure to move across the model hierarchy that allows understanding how changes in biological parameters - physiological, pathophysiological, or because of new data - impact the behaviour of the network. We apply this approach to mammalian primary visual cortex, and examine how the different models in the hierarchy reproduce functional signatures of this area, in particular the tuning of neurons to the orientation of a visual stimulus. Our work provides a navigation of the complex parameter space of neural network models faithful to biology, as well as highlighting how simplifications made for mathematical convenience can fundamentally change their behaviour.

## 1 Introduction

Theoretical modelling of a neural system can be accomplished with different degrees of biological detail and, conversely, different degrees of mathematical abstraction. Arguably, including more biological detail based on experimental data provides a model which is somehow a more faithful surrogate for the original system. But because detail does not necessarily yield understanding, a more abstract description with fewer parameters and thus fewer degrees of freedom offers the possibility of better insight. It follows that mapping between different categories of models is necessary to relate analytical results obtained from more abstract models, to simulations from detailed models which can hardly be analyzed in their parameter space. Eventually, this sort of back and forth process can facilitate understanding about how specific model assumptions may be linked to detailed model behaviour.

In this paper we address the challenge of parameter reduction and quantitative mapping from original biophysical quantities to simplified model parameters, using the orientation hyper-column, or pinwheel, architecture in the visual cortex of higher mammals as a model system. The orientation hyper-column is a functional unit providing organization of cellular tuning with respect to stimulus orientation, and many theoretical approaches have been used to study this neuronal network. At one extreme are simplified population models that retain only gross features of the modeled system, but allow for systematic analytical and numerical investigations, and thus a formal understanding of underlying mechanisms to the extent that the model is reliable. The canonical model in this sense is the firing-rate (FR) ring model with a single neural population type and a one dimensional ring architecture corresponding to preferred orientation [1] and [2]. At an increased level of complexity, [3] and [4] also consider populations based on rate models in a 2D distributed architecture corresponding to the cortical surface geometry. Extensions of population models have taken into account different connection profiles [4] or delays [5], both excitatory and inhibitory neuronal populations [3], or more elaborate rate models [6].

At the other extreme are biophysically-detailed models which attempt to incorporate known physiology and synaptic anatomy of the system, giving a coherent description that bridges established parameters and those which are unknown yet essential. For visual cortex there is a wealth of experimental data including various neuron types and their intrinsic biophysics, the functional architecture of thalamic input to the system, the kinetics of the synaptic conductances, and the schematic of the intra-cortical synaptic connections. Numerical simulations of such models, for example [7], [8], [9] and [10], have provided realistic reproductions of experimental data, and can provide explicit predictions for subsequent experimental studies. In general, a complex model can incorporate the available data at will, motivated by the fact that *a priori* we do not know which details are fundamental for function, e.g. the detailed dynamics of receptive fields.

Here we consider a hierarchy of single layer hyper-column models (Table 1 and Figure 1), at the top with a reasonably detailed model that considers biophysical membrane properties of cortical excitatory and inhibitory neurons with synaptic conductances described by second order kinetics. The synaptic architecture of this model arises from a 2D anatomical distribution of the two neuron types over the cortical surface, with spatial interactions defined by isotropic intracortical connections, and thalamic input according to an assumed pinwheel architecture. This full model has been realized with the help of the conductance-based refractory density approach [11]. Via step by step simplifications of the synaptic architecture and cellular elements, comprising five intermediate models, we arrive at the canonical firing rate ring model. We require a consistency between nearest-neighbors within the hierarchy of models to validate each mapping, in the sense that any qualitative or quantitative change in the behaviour of a given model must be linked to specific assumptions made at each stage of the reduction.

**Table 1.**
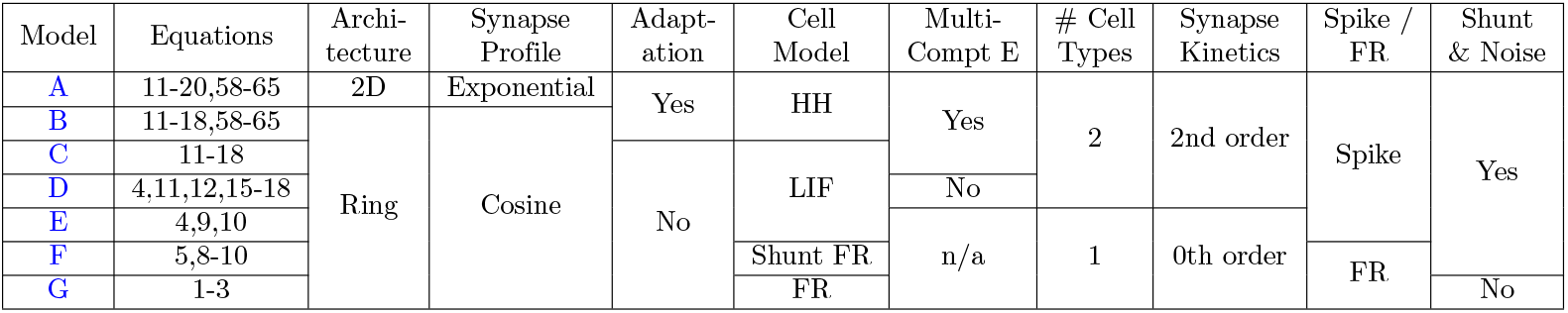
Summary of Network Models. Abbreviations: 2D, 2 dimensional; HH, Hodgkin-Huxley; LIF, leaky integrate-and-fire; FR, firing rate; Multi-Compt E, multiple compartment excitatory neuron

**Fig 1.**
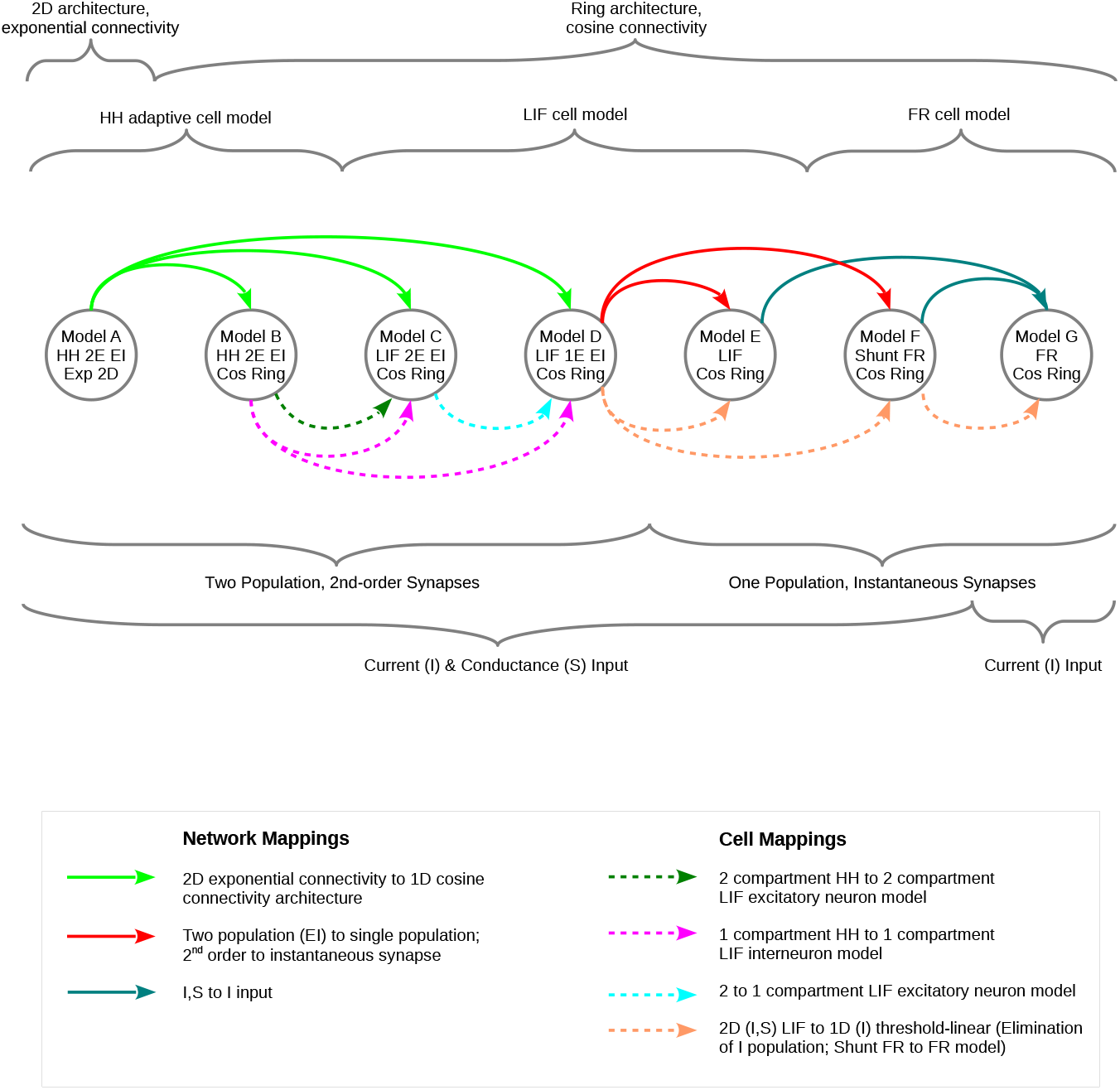
General characteristics of models, and the mappings between them, including steps for network parameters (solid arrows) and cellular parameters (dashed arrows). Arrows that arise from, or terminate at, multiple source or destination models, respectively, indicate a common characteristic of the models specific for a given mapping. For explanation of model abbreviations, see Table 1.

Note that this particular hierarchy is not comprehensive, e.g. an arbitrarily elaborate network architecture may be evaluated with a simple cell model, or vica-versa. Rather, the motivation for the models studied here is that several are directly linked to previous published results. Furthermore, the methodology that we develop may be readily adapted to other choices of hyper-column models, as well as network architectures of other cortical areas.

## 2 Results

We present the results as follows. We begin by describing the hierarchy of network models and the relevant cellular models from simplest to most complex, thus progressing from the abstract current-based firing rate ring model, until the biophysically detailed 2D cortical model that considers Hodgkin-Huxley membrane channels, synaptic currents and conductances. Next, we describe the mapping steps at the cellular and network levels, now in the opposite direction, from the most detailed network and cell models, until the firing rate ring model. Once the mapping strategy and formulas are described, we present an analysis of the parameters of the single population ring models, taking into account the significance of various biophysical values taken from experiments. Numerical simulations of the different network models are then presented, focusing on the similarities and differences in the response steady-state and dynamics to a stereotyped visual stimulus sequence.

### 2.1 Model hierarchy

We consider two geometries of the orientation hyper-column circuit in primary visual cortex, thus a 2-dimensional geometry that maps the cortical surface to function according to the orientation preference hyper-column map found in higher mammals, and a one dimensional ring geometry, where orientation preference corresponds to the position along the ring. Table 1 and Figure 1 present the model hierarchy, emphasizing the basic characteristics of the models in terms of network architecture and cellular properties. The models are assigned letters according to the presumed hierarchy, ranging from the most elaborate 2-dimensional, biophysical, model A at the top, to the simplest version of the ring model, model G, at the bottom. For clarity, in this section we present the models in order from the simplest to the more complex (model G, model F, model E, etc.).

#### 2.1.1 Current-based firing rate (FR) ring model with instantaneous synapses (Model G)

The simplest network that we consider is the classical FR ring model described by [1] (see also [2]). The network consists of a ring in orientation space, parameterized by *θ*, of a single population of firing rate neurons whose activity is given by the spike rate *ν*(*t,θ*):

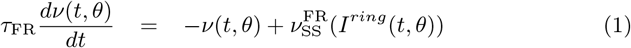

where *τ*_FR_ is a phenomenological time constant of the rate relaxation. The steady-state firing rate of the population, 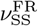, is given by a threshold-linear transfer function:

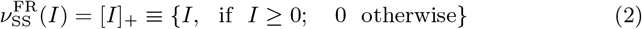

The network architecture is fully defined by the expression for the input *I^ring^*(*t,θ*):

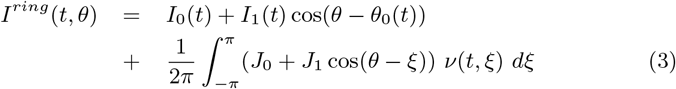

where *I*_0_ and *I*_1_ are amplitudes of non-tuned (stimulus background) and tuned thalamic input, respectively, and *θ*_0_ is the orientation of the input; *J*_0_ and *J*_1_ are amplitudes of non-tuned and tuned recurrent intracortical input, respectively. These parameters aggregate the effects of spatial connections, synaptic strengths and dynamics between populations of different types. In general the thalamic input is a function of time, thus *I*_0_(*t*), *I*_1_(*t*) and *θ*_0_(*t*). Note that this model incorporates the simplest synapse model, where input current arising from intracortical activity is directly proportional, and thus instantaneous, with respect to the firing rates of the pre-synaptic population (which also allows the thalamic firing rates to be implicit in this equation).

In this single population model the effect of inhibitory versus excitatory cortical neurons is implicit. The original interpretation of *J*_0_ and *J*_1_ with respect to excitation and inhibition is given in [1]: “The constant *J*_0_ represents a uniform all-to-all inhibition; *J*_1_ measures the amplitude of the orientation-specific part of the interaction. Neurons with similar preferred orientations are more strongly coupled excitatorally than neurons with dissimilar ones.” As originally formulated by [1] (see also [2]), the values of *I*_0_ and *I*_1_ were chosen so that the feedforward input is non-negative, since there is no experimental evidence for inhibitory thalamocortical pathways [12]. In Section 2.2.4 we will present an alternative interpretation of these coefficients that arises from the mapping, specifically that takes into account the implicit inhibitory cortical population of this model.

#### 2.1.2 Conductance-based shunt FR and leaky integrate and fire (LIF) ring models with instantaneous synapses (Models E, F)

We now elaborate the single population current-based ring by incorporating synaptic conductances as well as currents, and by including a membrane current noise term. We consider firing rate and spiking versions of the ring, based on a shunt firing rate neuron (model F), and a leaky integrate and fire neuron (model E), respectively.

The conductance-based LIF ring model E explicitly considers the effect of spikes on the membrane potential, *V*. The current equation for a single compartment LIF neuron at location *θ* along the ring is given by:

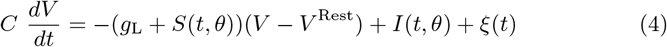

where *C* is the cell capacitance, and *g*_L_ is the leak conductance. Synaptic input is defined as the total synaptic conductance, *S*(*t,θ*), and the total synaptic current, *I*(*t, θ*), the latter measured with the cell held at the resting potential (note that a current term representing external stimulation may be included without loss of generality). *ξ*(*t*) is a Gaussian white current noise process characterized by its mean, 〈*ξ*(*t*)〉 = 0, and auto-correlation, 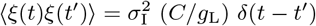, and *σ*_I_ is the noise amplitude. When *V* crosses a defined threshold voltage *V*^Th^, the neuron ”fires” and the membrane voltage is immediately reset to a defined voltage *V*^Reset^. For simplicity, in particular to allow a direct mapping to a firing rate model, we neglect an absolute refractory period under the assumption that the evoked firing rates are well below maximal.

We note that a noisy population of LIF neurons with instantaneous synapses can be evaluated either with Monte-Carlo methods, or with a probabilistic approach based on the Komogorov-Fokker-Planck (KFP) equation. We use the latter approach, described in the Methods (Section 4.1), to derive the simulations presented in the Results. Thus, in the presentation below of the input terms for model E we will refer to specific formulae of the KFP approach.

The average steady-state firing rate of the LIF neuron defined by Eq 4 provides a hybrid “shunt FR” network model (model F), situated between the previously described current-based FR model G, and the conductance-based LIF model E, that takes into account noise as well as the impact of synaptic input on the membrane time constant. For a given input *I* and *S*, the average steady-state firing rate of the LIF model, 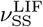, is given analytically by [13] (see also Figure 14):

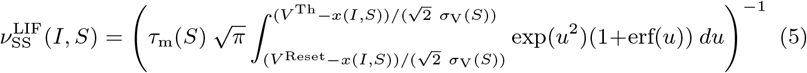

where the asymptotic potential, *x*(*I, S*), the steady-state voltage dispersion, *σ*_V_(*S*), and the effective membrane time constant of the LIF neuron, *τ*_m_(*S*), all depend on the synaptic conductance *S*:

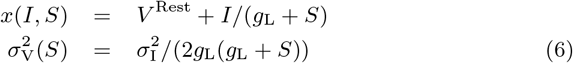

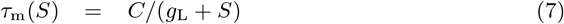

The shunt FR model F is then defined by using Eq 1 of the current-based FR model G, now with the steady-state solution given by 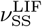 of the LIF model E:

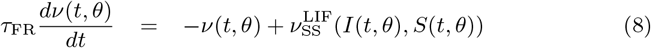

The form of the synaptic current and conductance network connections of models E and F are identical; we assume they recapitulate the constant plus cosine form of the current-based FR ring model G (Eq 3) and behave identically as a function of *θ*:

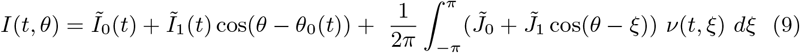

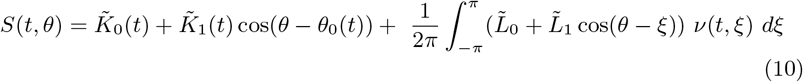

Thus, these expressions supply the *I* and *S* terms of Eq 8 for the shunt FR ring model F, and likewise the *I* and *S* terms for Eq 4 for the single population LIF ring model E (specifically, for Eqs 53 and 54 of the KFP evaluation of an infinite LIF population, Section 4.1). As with the current-based FR ring model, synapses are instantaneous for models F and E, in this case with the intracortical synaptic current and conductance input terms being directly proportional to the pre-synaptic firing rate *ν*(*t, ξ*), and as before with thalamic firing rates implicit in these equations. The population firing rate, *ν*(*t,θ*), for the shunt FR model is given directly by Eq 8; for LIF neurons (also the HH neurons discussed below), *ν*(*t,θ*) is defined by summing all spikes in the population at location *θ*, *n*_act_, over a time window Δ*t*, dividing by the number of neurons, *N*, and Δ*t*, as Δ*t* goes to zero:

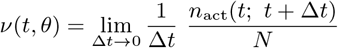

#### 2.1.3 Two population LIF and Hodgkin-Huxley (HH) ring models with the second order synapse kinetics (Models B, C, D)

We next consider two population ring models that include explicit excitatory (E) and inhibitory (I) neuron populations, based either on the LIF cell model, or more complex Hodgkin-Huxley (HH) type cell models. The two population LIF-based ring models include a single compartment inhibitory cell model, with either a one compartment LIF excitatory cell model (model D), or a two compartment LIF excitatory cell model (model C). Note that by definition, the standard LIF model as used here does not exhibit adaptation. At a next level beyond the LIF description, HH-type models are the standard framework for describing cellular dynamics tied to explicit biophysical mechanisms, including voltage-dependent membrane channels [14]. The HH-based ring model (model B) includes an adaptive two compartment excitatory cell model and a non-adaptive single compartment inhibitory cell model. The intracortical weighting functions for the excitatory and inhibitory pathways in all three of these two population ring models, fitted to anatomical data, follow the constant plus cosine form presented above.

The two population models allow distinct cellular and synaptic properties for excitation and inhibition, which in turn motivates the consideration of more elaborate synaptic kinetics. Each cortical population, therefore, has three types of synapses: those arising from thalamic excitatory (Θ), from intracortical excitatory (E), and from intracortical inhibitory (I) pathways. Accordingly, each connection in the network is denoted by a double index *ij* where *i* and *j* indicate the pre- and post-synaptic population, respectively. The complete synaptic input to a target population of type *j* is then given by the total synaptic current measured at rest, *I_ij_*, and the total synaptic conductance, *S_ij_*, summed over the different synapse types arising from pre-synaptic populations *i*.

The excitatory and inhibitory synaptic currents measured at rest for each cell type *j* at location *θ*, and the associated synaptic conductances, are thus:

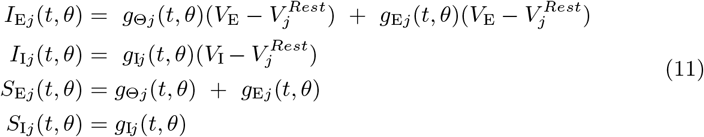

where *V*_E_ and *V*_I_ are the excitatory and inhibitory reversal potentials, and 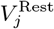 is the resting potential of the target population.

For the one compartment LIF and HH cell models (both cell types in model D; the inhibitory cell type in models B and C), we refer to the total input, thus:

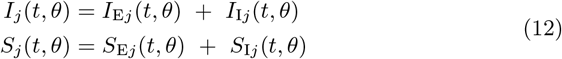

The membrane current equation for the single compartment LIF and HH models of cell type *j* at location *θ* is then:

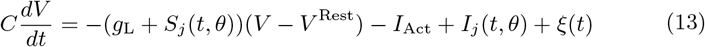

where *I*_Act_ is the current via active, voltage-gated channels for HH neurons, and ignored for LIF cells.

For the two compartment HH and LIF excitatory cells of models B and C, the post-synaptic sites are pathway specific. Thus we refer directly to Eqs 11 for the excitatory input exclusively on the dendrite, *I*_EE_(*t, θ*) and *S*_EE_(*t, θ*), and for the inhibitory input exclusively on the soma, *I*_IE_(*t*, *θ*) and *S*_IE_(*t*, *θ*). The two compartment models are an approximate solution of two boundary problems for a spatially distributed passive dendrite and a point passive soma, following the description in [15] (see also [9] and [16]). The problems correspond to the two cases of, first, dendritic synaptic current estimation under somatic voltage clamp (the reverse voltage-clamp problem) and, second, somatic voltage measurement under current-clamp in response to the previous estimates of the dendritic synaptic current.

The membrane current equations for the soma voltage *V*(*t*) and the dendrite voltage *V*_D_(*t*) for this excitatory cell model at location *θ* are:

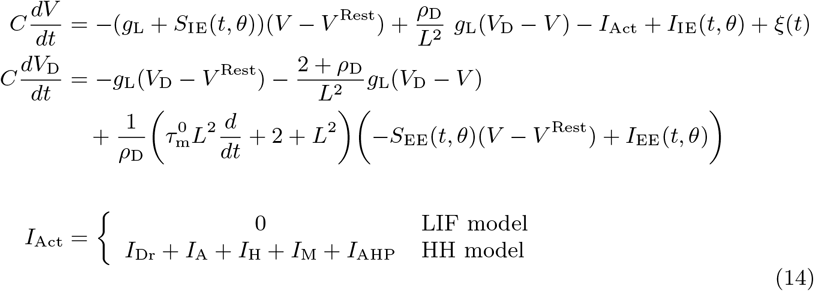

where 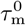 is the resting time constant (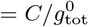, where 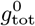 is the total conductance at rest), *ρ*_D_ is the ratio of the dendritic to somatic passive membrane conductances, and *L* is the dendritic length in units of the characteristic length (see Methods, Section 4.3).

Notably, spike generation in the HH cell models used here is governed by an explicit threshold mechanism, reminiscent of the LIF model, that replaces the spike generating current *I_Na_*. This modification, described in detail in the Methods (Section 4.2), allows for efficient evaluation of neuron populations that still considers the impact of the various HH currents that comprise *I*_Act_ (Eqs 13 and 14). It is important to note that this approach was taken for its computational advantage alone; using a standard HH cell model in the framework of the network simulations will give very similar results [17] (see also Section 4.2).

The connectivity rules for all the two population models are written in terms of pre-synaptic firing rates, *φ_ij_*, to facilitate incorporating synaptic kinetics in the translation of pre-synaptic activity to the post-synaptic conductances *g_ij_*(*t*) that figure in Eqs 11. The relation between *φ_ij_*(*t*) and *g_ij_*(*t*) is given by a second order kinetic model that implicitly accounts for mechanisms such as axonal delays, receptor dynamics and dendritic integration:

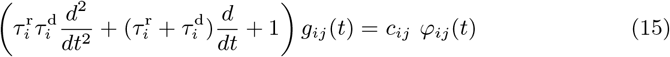

with the scaling “synaptic capacity” *c_ij_* described in Section 4.5. The thalamic inputs driving *g*_Θ*j*_(*t*, *θ*) of the excitatory and inhibitory cortical populations are taken to be identical, and follow the constant plus cosine form of the single population ring models:

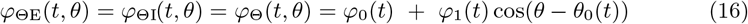

where *φ*_0_ is the un-tuned stimulation background, *φ*_1_ is the tuned part of the cortical input, and *θ*_0_ is the stimulus orientation angle. As mentioned, the thalamocortical pathway is exclusively excitatory.

Intra-cortical connections driving *g_ij_*(*t*, *θ*) of the two populations are given by:

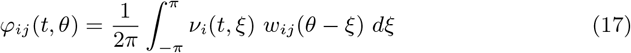

where the weight function, *w_ij_*(*ϕ*), again recapitulates the constant plus cosine form of the single population models, with the parameter *q_ij_* to take into account the orientation tuning between different types of neurons:

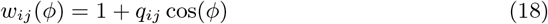

Note that, implicitly, 0 ≤ *q_ij_* ≤ 1. This approximation will be developed in Section 2.2.1.

#### 2.1.4 Two population, two dimensional HH model with the second order synapse kinetics (Model A)

As the most elaborate model, we consider an anatomical 2D cortical network of the two compartment excitatory, and the single compartment inhibitory, HH cell models developed in the previous section. As for the two population ring models, the current and conductance inputs are driven by pre-synaptic firing rates.

As opposed to the angular functional index *θ* of the ring models, in the 2D model each population is parameterized by its position on a cortical surface map of adjacent, radially-symmetric hyper-column pinwheels [3] (Figure 2). The 2D architecture is captured entirely by new expressions for the firing rates *φ*_Θ_ and *φ_ij_* that are now functions of the *x* and *y* coordinates of the target cells. Thus, the inputs of the 2D model recapitulate Eqs 11–15, with arguments (*t, θ*) of the ring models replaced by (*t, x, y*).

**Fig 2.**
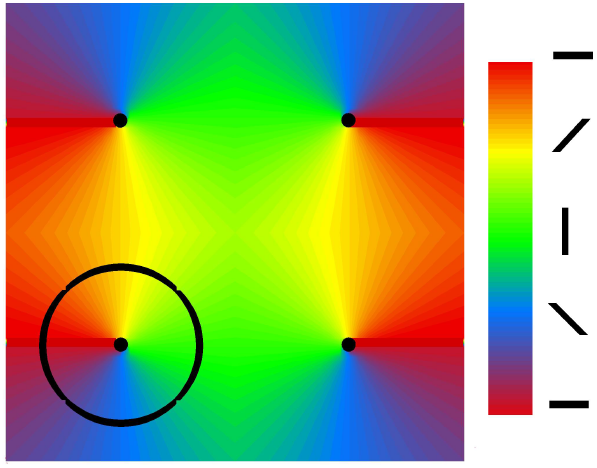
Cortical domain containing 4 hypercolumns that provides the reference geometry for the 2D model A; the dots mark the pinwheel centers; the colors mark the orientations coded by bars in the legend. The circle at the lower left quadrant corresponds to the activity profile of this model shown in the first panel of Figure 8.

An orientation column (Figure 2) corresponds to a hyper-column sector, containing populations with similar preferred orientations at varying distances from the hyper-column center. The functional architecture of each hyper-column pinwheel arises from a thalamic input whose orientation tuning corresponds to the angle of the target location with respect to the center of its hyper-column [8], [3]. Hyper-columns are arranged so that those with a clockwise progression of orientation columns are adjacent to those with counterclockwise progression, thus adjacent orientation columns from different pinwheels have the same orientation preference. The centers of the hyper-column pinwheels are distributed on a rectangular grid with pinwheel radius *R* and indexed by *i_PW_* and *j_PW_*. Here we consider a square containing 4 pinwheels on the cortical surface. The coordinates of the pinwheel-centers are *x_PW_* = (2*i_PW_* − 1)*R, y_PW_* = (2*j_PW_* − 1)*R*. The orientation angle for the point (*x,y*) which belongs to the pinwheel (*i_PW_, j_PW_*) is defined as *θ* = arctan((*y* − *y_PW_*)/(*x* − *x_PW_*)). The progression is determined by the factor (−1)^*i_PW_*+*j_PW_*^.

Thus, the thalamic input in terms of effective firing rate is described with an elaboration of Eq 16:

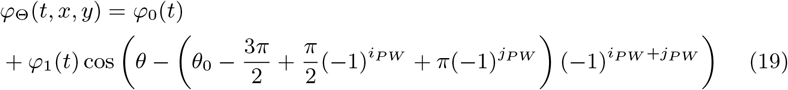

It is important to note that, as with the ring models, this constant plus cosine form of the thalamic input constitutes an implicit model of the retino-geniculo-cortical pathway; this is equivalent to each cortical neuron receiving an oriented receptive field corresponding to a single sub-region of a classical simple cell. The orientation dependence of the input in particular characterizes the degree of symmetry breaking at the input stage, parameterized here by *φ*_1_ and *φ*_0_. The homogeneous case, where *φ*_1_ = 0, represents thalamic input with no dependence on stimulus orientation. The maximum selectivity of this model for the input stage, where *φ*_1_ = *φ*_0_, is a pure shifted cosine dependence.

Intra-cortical connections between populations are local and isotropic with an exponential decay [12], with the effective pre-synaptic firing rate *φ_ij_*(*t, x, y*) given by:

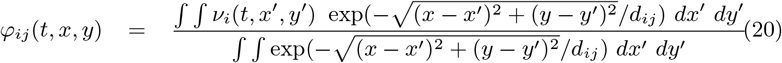

where, as above, *i* and *j* correspond to the cell type (E or I), *ν_i_*(*t, x′, y′*) is the firing rate of pre-synaptic cortical population i at location (*x*′, *y*′), and *d_ij_* is the characteristic anatomic length of the *i* to *j* pathway. Note that in contrast to the explicit tuning of intracortical connections in the ring models (parameterized by *θ*), these connections have an implicit functional orientation tuning since neighboring orientation columns receive similar oriented thalamic inputs.

### 2.2 Mapping of network and cellular parameters

We now describe the quantitative mappings between the various models at the network and cellular levels, which allow a path from the parameters of the full HH EI 2D model A, to the different LIF networks models, and finally to the current-based FR ring model G. In contrast to the previous section, we present the mappings from the more complex to the less complex models. The essential goal of these cellular and network mapping steps, as diagrammed in Figure 1, is to systematically map the high-dimensional full model, with multiple non-linearities, to a low-dimensional model with a single threshold non-linearity.

#### 2.2.1 Network mapping: 2D cortical architecture to the cosine ring models

The first mapping step at the network level is to reduce the 2D geometry of the pinwheel architecture in anatomic space to a 1D ring geometry in orientation space (solid green arrows in Figure 1). We consider the synaptic strength function for the intracortical connection between two points 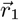 and 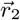 on the cortical surface (corresponding to the locations (*x, y*) and (*x′, y′*) in Eq 19):

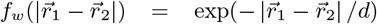

where *d* is the characteristic length of the connection. The weight of the connections of two elementary areas Δ*S*_1_ = *r*_1_ Δ*θ*Δ*r*_1_ and Δ*S*_2_ = *r*_2_Δ*ξ*Δ*r*_2_ at the points shown in Figure 3A is:

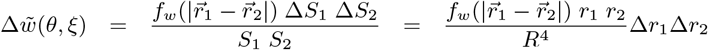

where 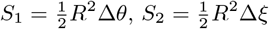 are the areas of the sectors corresponding to two radial vectors emanating from the center of a pinwheel with radius *R* and having the orientation angles *θ* and *ξ* (Figure 3A). Combining the last two expressions, we integrate over the vectors to obtain the total connection weight 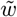:

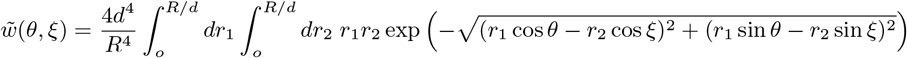

**Fig 3.**
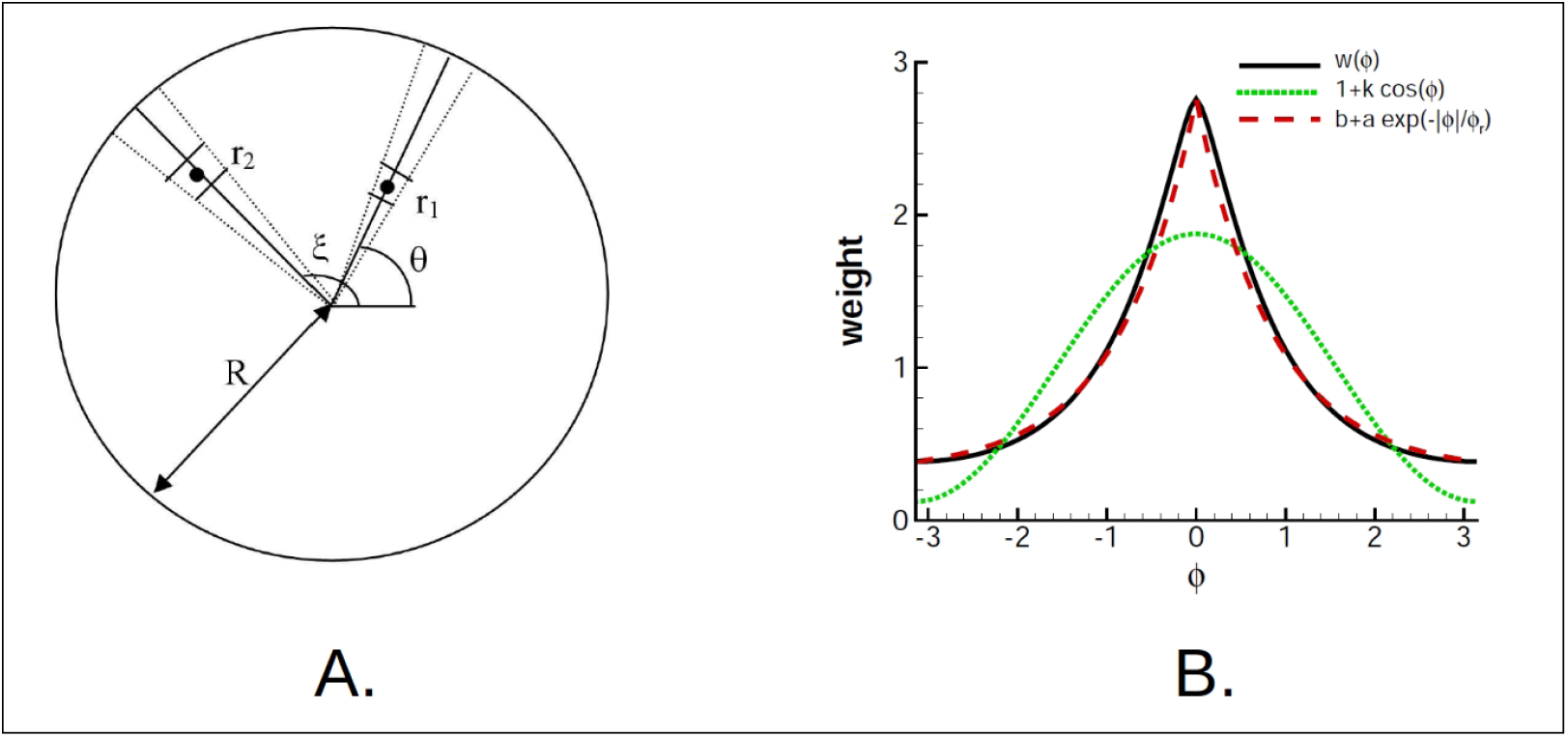
A. Reduction of 2D pinwheel geometry to the ring-geometry requires integration over the radial vectors with the angle *ξ* − *θ* between them. B. Mapping 2D pinwheel geometry to the ring, showing the weight function *w*_0_(*ϕ*) according to the Eqs 21 and 22 (solid line), and its approximations of the forms 1 + *k* cos(*ϕ*) (dotted line) and *b* + *α* exp(−|*ϕ*|/*ϕ_r_*) (dashed line) for the case of *R* = *d*/2.

We now find 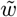 for the angle difference *α* = *ξ* − *θ*. By rotating the coordinates such that *θ* = 0 and *α* = *ξ*, we obtain:

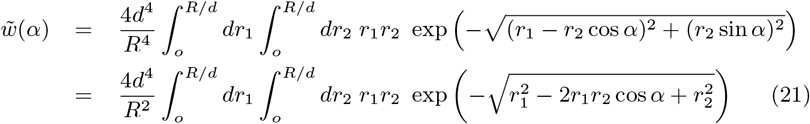

The resulting normalized weight

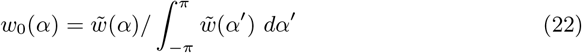

is shown in Figure 3B by the black curve. As mentioned in Section 2.1.3, we approximate *w*_0_(*α*) with a constant plus cosine form for models B, C and D (Eq 18, parameterized by *q_ij_*; reference Figure 3B, dotted line). We also considered an exponential form *w_ij_*(*ϕ*) = 1 + *q_ij_* exp(|*ϕ*|/*ϕ_ij_*) (Figure 3B, dashed line), which gives a more precise mapping of the anatomically based 2D cortical geometry to the ring [18], which we will discuss later.

#### 2.2.2 Cellular mappings: HH to LIF neurons, and 2 compartment to 1 compartment LIF excitatory neuron

The mappings of the HH neuron models to their respective LIF models were made as follows. For the population of one compartment HH inhibitory neurons, the steady-state transfer function between current input and firing rate (f/I) of a one compartment LIF model, with the same passive parameters of the HH neuron, was fitted to the f/I curve of the HH model by adjusting the LIF reset potential, while retaining the threshold potential *V*^Th^ of the HH inhibitory model (dashed purple arrows in Figure 1). A similar fitting procedure was done for the two compartment HH excitatory neurons, but importantly using a non-adapting version of this model. Thus, as a first step, the two adaptation currents of the excitatory neuron, *I*_M_ (intermediate time-scale voltage-dependent current) and *I*_AHP_ (slow voltage and calcium-dependent current), were blocked, and the f/I was obtained. The f/I curve of a two compartment LIF model was then fitted to the non-adapting HH f/I curve, with the same passive parameters and threshold potential *V*^Th^ of the HH excitatory model (dashed green arrow in Figure 1). The comparison of the spike timings in response to constant input is shown in Figure 4.

**Fig 4.**
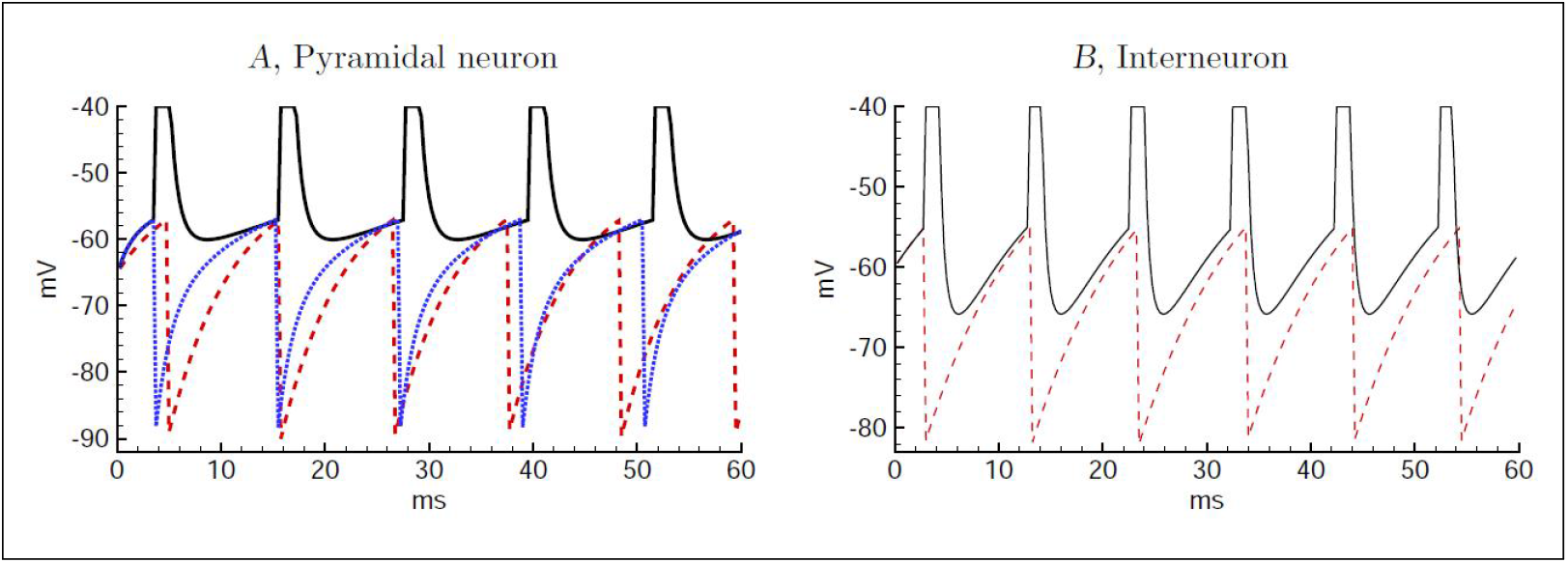
Comparative spike train responses of the mapped single neuron models: HH tc LIF, and 2 to 1 compartment LIF. A) The spike train for the non-adaptive 2-compartmental HH pyramidal neuron (solid line), the 2 compartment LIF model (dashed line) and the 1 compartment LIF model (dotted line), the latter two with 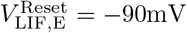. The input to the dendrite of the two compartment models was *I* = 1.2nA and *S* = 0.035 *μ*S; according to the ratio of the dendritic and somatic input conductances, *G*_in,d_/*G*_in_, the current and conductance input to the 1-compartmental neuron was adjusted to 0.62nA and 0.018*μ*S, respectively. B) The spike train for the HH interneuron (solid line) and the corresponding LIF model with 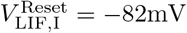 (dashed line), in response to *I* = 0.2nA and *S* = 0.

To map the two compartment LIF excitatory cell model to single compartment LIF model (dashed cyan arrow in Figure 1), we fixed the passive input conductance of the latter to the somatic input conductance of the former, and decreased the membrane time constant from 14.4 to 10.3 ms, reflecting the fact that the effective time constant of a two compartment model is smaller than intrinsic membrane *τ*_m_. Because of different expressions for the input conductance in two and single compartment models, expressed by the ratio of the dendritic and somatic input conductances, *ρ*_D_ = *G*_in,d_/*G*_in_ (Eqs 63 in Methods, Section 4.3), the capacitance of the one compartment model changed slightly from 0.25 to 0.27nF. The synaptic input to the dendrite of the two compartment model (excitatory inputs from the thalamus and the excitatory cortical population) were adjusted for the one compartment model by *ρ*_D_ so that the sizes of the synaptic conductances relative to the post-synaptic input conductance were maintained (ref. Figure 4A for a comparison of the spike times of the two models).

#### 2.2.3 Cellular mapping: 2-dimensional LIF steady-state to 1-dimensional threshold-linear f/I relation

We now describe the mapping of the steady-state expression for the current (*I*) and conductance (*S*) driven LIF model, 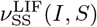 (Eq 5), to a one dimensional threshold-linear model, thus with the current and conductance inputs combined *a priori* (dashed orange arrows in Figure 1). This transformation is used twice. First, this is used for eliminating an explicit inhibitory population when going from the two population LIF ring model D, to the conductance-based single population models E and F (Section 2.2.4). Second, this step is used for mapping from the conductance-based shunt FR model F, to the classical current-based FR model G (Section 2.2.5).

We first assume that the square of the noise current amplitude, 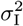, is marginally proportional to the synaptic conductance:

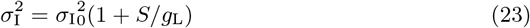

with the noise level at rest *σ*_I_0__. This approximation reflects the fact that the synaptic current fluctuations typically increase as more synaptic channels are activated. The ionic channels provide independent additive stochastic currents, thus the variation of the total current fluctuations is the sum of variations of the currents from separate channels, which is roughly proportional to the number of activated channels. The later is characterised by the total conductance. Importantly, for the LIF neuron this leads to constant voltage dispersion due to noise, *σ*_V_(*S*) (Eq 6):

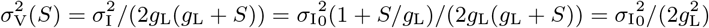

Independence of the voltage dispersion on the total conductance corresponds to the case of thermal noise in RC circuits [19]. The scaling Eq 23 and, consequently, the constant *σ*_V_ were derived in the particular case of a balanced excitatory and inhibitory synaptic input by Chance et al. [20].

Given the form of equation (5), this in turn allows a recasting of the steady-state solution 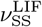 as a one dimensional function:

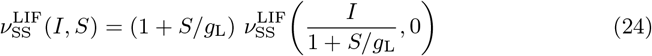

We note that this reduction of a 2 to 1 dimensional solution is unique for this noise model, and is an important analytical property of the noisy LIF model. In the notations from [20], the formula of Eq 24 corresponds to the relationship *r* = *g f* (*I/g*) with the total conductance *g*, the input current *I* and the firing rate *r*. With respect to the modulation of the f/I curve gain, we see that the shunt *g* provides a divisive effect on the gain *f*′ (*I/g*) only if the second derivative of *f* is positive.

Next, we approximate 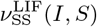 for *S* = 0 as a one dimensional threshold-linear function, parameterized by the slope *k*^LIF^ and the rheobase current 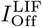:

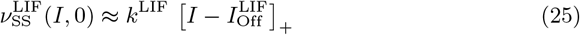

The approximation parameters for the LIF models introduced in the previous Section 2.2.2, thus 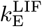 and 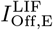 for the excitatory LIF neuron, 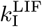 and 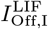 for the inhibitory LIF neuron, are obtained by fitting to 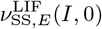 and 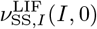, respectively. The precision of this approximation for the excitatory cell type can be seen from Figure 5, which compares the threshold-linear approximations for noisy LIF and HH cell models plotted for parameters given in Section 2.3.

**Fig 5.**
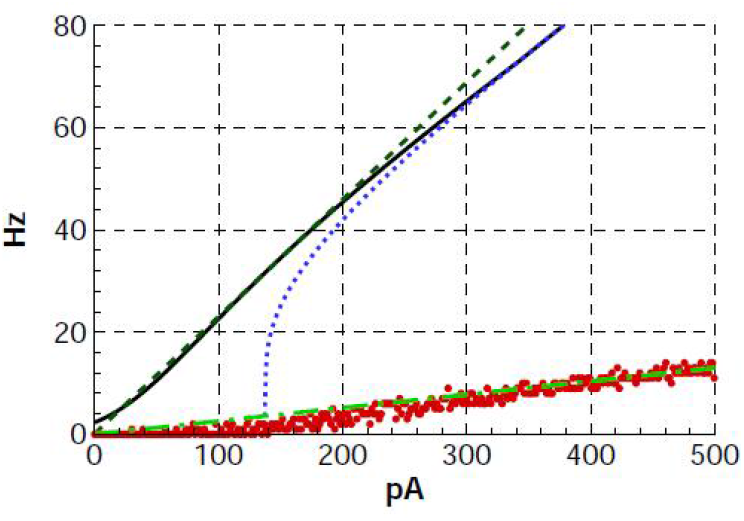
Comparisons of threshold-linear approximations for noisy LIF and HH excitatory cell models. Threshold-linear approximation (dashed line) of the LIF noisy neuron firing rate (Eq 5, solid line). To compare, the response of a LIF neuron with no noise is shown by the dotted line. The steady-state rate of the adaptive neuron described in Section 4.3 is shown by red dots, each dot corresponding to one stimulation current amplitude and being obtained by Monte-Carlo simulation with explicit noise during 2 seconds. The dashed-dotted line is the threshold-linear approximation with a gain of 0.026 Hz/pA.

Combining the last two expressions obtains an approximate one dimensional threshold-linear dependence of the rate on *I* and *S*:

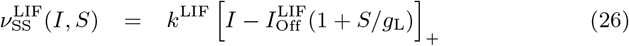

Thus, the effect of the synaptic conductance is to modulate the threshold current of the LIF neuron 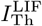:

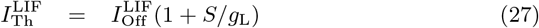

A similar approximation was described in [6], in the context of fitting threshold-linear functions to the numerically obtained steady-state transfer function of a HH neuron with a slow potassium A-current, specifically approximating the impact of the cell conductance as a rightward shift in the transfer function. Brizzi et.al. [21] showed that this approximation was justified for spinal motor neurons *in vivo*, and Persi et al [22] generalized the approach for HH neuron models with current noise. The difference with our interpretation arises from the application of the similarity law (Eq 24), which is exact for the particular noise model (Eq 23) and the LIF neuron. Note, in the aspect of the f/I curve gain problem, we see from Eq 26 that the channel-to-channel independent noise results in shunt-independent gain of f/I curve, i.e. the shunt provides a pure subtractive effect.

As shown in the Section 2.3, the quality of these approximations allows to obtain consistent mapping of the model parameters. Note that the choice of a threshold-linear transfer function approximation for the steady-state is not essential for the mapping from the two population to one-population models, nor between the conductance-based to current-based FR models, and thus a more elaborate one dimensional function may be used. However, the threshold-linear approximation not only simplifies these mappings (e.g. solving for *ν*_I_ in Section 2.2.4), but it is also necessary for the final mapping to the classical threshold-linear FR model that is especially amenable to analytical results.

Considering the form of the synaptic current and conductance introduced previously, we can simplify the mapping further. Now we explicitly indicate the LIF cell type *j*. First, we express the threshold voltage of the LIF neuron in terms of its rheobase current 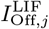 and input conductance *g*_L, *j*_:

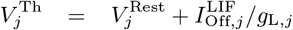

We now define 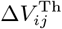 as the driving force relative to the target’s threshold potential:

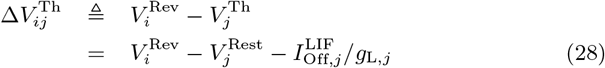

where 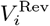 is the reversal potential of synapse type *i*. With this definition of 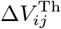, we introduce the network input, recalling the definitions of *I_j_* and *S_j_* (Eqs 11–12), and finally rewrite Eq 26 for 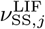:

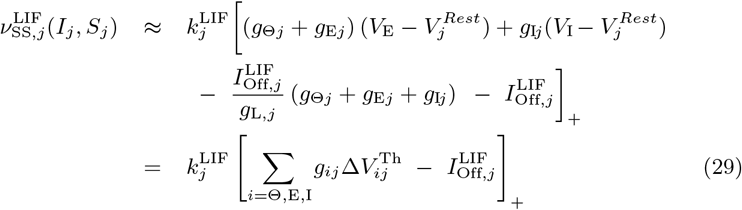

#### 2.2.4 Network mapping: Conductance-based two population ring to single population ring

We now map the current and conductance inputs *I_j_*(*t, θ*) and *S_j_*(*t, θ*) (Eqs 12) of the two population ring model D, to the *I*(*t, θ*) and *S*(*t, θ*) inputs (Eqs 9,10) of the single population LIF and FR with shunt ring models (models E and F; ref. solid red arrows in Figure 1). This mapping effectively eliminates the explicit inhibitory population, replacing it by an instantaneous linear function of the excitatory activity. Note that the cell models in models E and F are derived from the single compartment excitatory LIF model of model D.

The first step is to replace the 2*^nd^*-order synaptic kinetics by instantaneous transfer functions, thus synaptic currents and conductances being directly proportional to the effective pre-synaptic rates, according to the steady-state solution for Eq 15, which is:

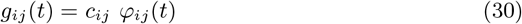

Now, the time dependence of the input is implicit:

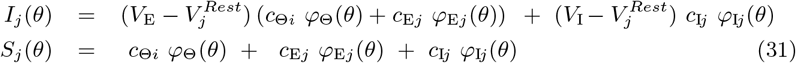

Next, we expand Eq 31 for the current input to excitatory neurons, *I*_E_(*θ*), using the pre-synaptic firing rates of the thalamic (*φ*_Θ_, Eq 16) and intracortical (*φ*_EE_ and *φ*_IE_, Eq 17) populations:

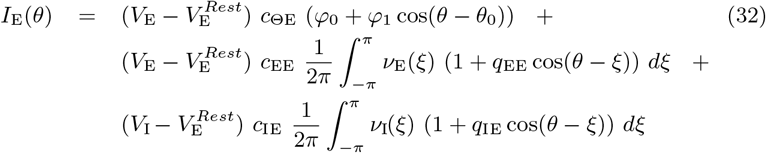

We now replace the inhibitory population rate *ν*_I_ in this equation by the steady-state equation for the inhibitory LIF model expressed as a linear function of the excitatory population rate *ν*_E_, based on three assumptions. First, we assume that the connections between inhibitory neurons in the cortex are local, thus:

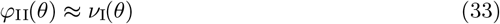

Next, we ignore the intrinsic dynamics of the inhibitory population and consider only its steady-state response, thus 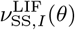, furthermore with the assumption that, on average, the inhibitory neurons around the ring are above threshold during stimulus presentation. Using the one dimensional approximation given by Eq 29 for the LIF inhibitory neuron (importantly neglecting the rectification operation because of the above-threshold assumption), Eq 31 for the current term (subsituting 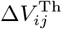 according to Eq 29), and the assumption of inhibitory-inhibitory locality (Eq 33), we obtain:

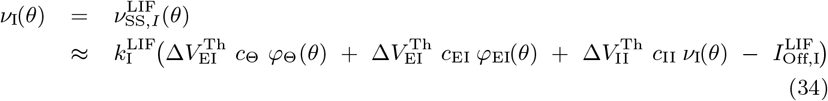

Eq.(34) may then be solved for *ν*_I_(*θ*):

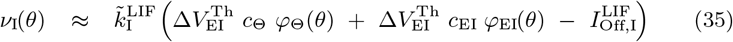

where:

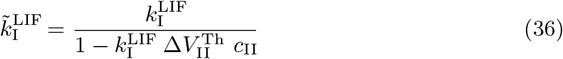

The term 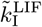 represents the reduction in the effective gain of the inhibitory population because of self-feedback, as such parameterized by the inhibitory-inhibitory pathway parameter *c*_II_. Assuming that the reversal potential of the inhibitory synapses is beneath the spike threshold, we note that 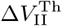 (Eq 28) must be negative, and therefore 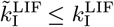.

Finally, we need to express the effective pre-synaptic rate of the excitatory input to the inhibitory population, *φ*_EI_(*θ*), in Eq 35 as a linear function of *ν*_E_(*θ*). For this we assume that the excitatory activity over the ring can be reasonably approximated by a shifted cosine centered at the stimulus orientation *θ*_0_:

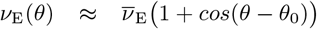

where 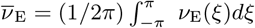, thus *ν*_E_(*θ*) averaged over the ring. Taking into account that:

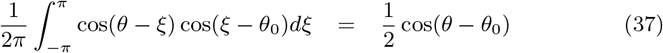

we obtain the following approximation for *φ*_EI_:

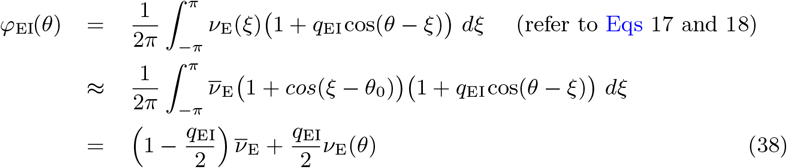

We can now re-write Eq 32 for the current *I*_E_(*θ*) as a function of only the excitatory population activity *ν*_E_(*θ*), substituting Eqs 16 and 38 for *φ*_Θ_ and *φ*_EI_, respectively:

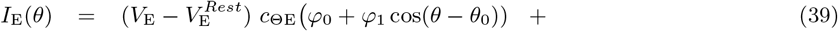

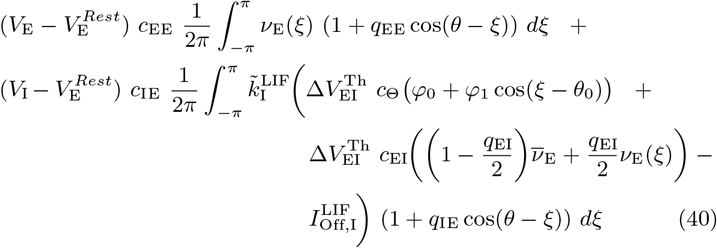

Again using the relation of Eq 37, we obtain the coefficients 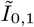 and 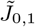 by comparing Eq 39 with the expression for *I*(*θ*) (ref. Eqs 9, i.e. setting *I*(*θ*) = *I*_E_(*θ*), with implicit time dependence):

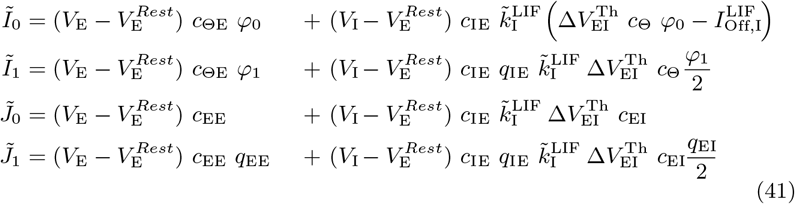

These equations provide the current terms in Eq 9, with the corresponding conductance terms in Eq 10 given by:

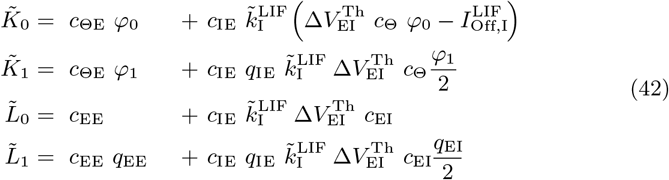

Note that the *C*_IE_ term in the above expression for 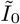 is not, in fact, driven by thalamic input (i.e. the thalamic firing rate term *φ*_0_), but rather is constant. Nevertheless, the formats of the defining equations for the inputs (Eqs 9 and 10) that distinguish between terms integrated with the network activity, from those that are “external”, oblige the inclusion of the *C*_IE_ in the “thalamic” component. In comparison to the thalamic terms, the input “driven” by intracortical activity is proportional to both 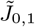 and 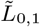 i.e. there is no non-driven, constant component.

To conclude, Eqs 41 and 42 constitute the mapping between the conductance-based two population ring models with 2*^nd^*-order synapses (models B, C and D) to the conductance-based single population ring models with instantaneous synapses (LIF model E, and shunt FR model F).

#### 2.2.5 Network mapping: Conductance-based single population ring to current-based FR ring

The final mapping is between the current and conductance inputs *I*(*t, θ*) and *S*(*t, θ*) of the conductance-based ring models E and F, to the current input *I*(*t, θ*) (Eq 3) of the current-based FR ring model G (dashed blue arrows in Figure 1)

As before we exploit the mapping of the two-dimensional steady-state expression for the LIF neuron, 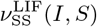, to the one dimensional threshold-linear approximation (Eq 26). Here, this step allows the elimination of the explicit conductance input, therefore to obtain the stationary solution 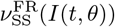 of the current-based FR model (Eq 1). As mentioned earlier, the conductance-based single population ring models reference the single compartment excitatory LIF model parameterized by 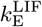 and 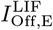 (Section 2.2). Thus, recalling the expression for 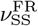 given by Eq 26, and expanding *I*(*t, θ*) and *S*(*t, θ*) (ref. Eqs 9 and 10) we obtain

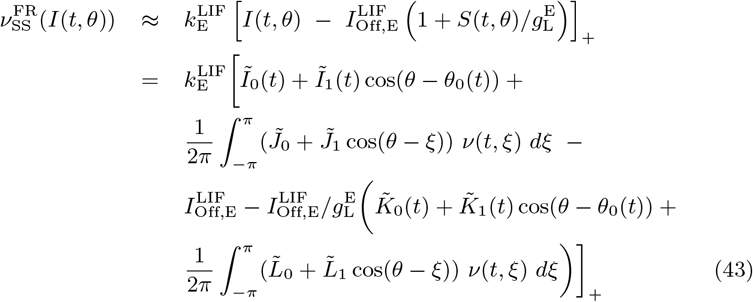

Note that, unlike the previous use of the one-dimensional transfer function approximation for eliminating an explicit inhibitory population, here there is no assumption that the input is suprathreshold. Comparing Eq 3 for *I*(*t, θ*) with Eq 43, we obtain:

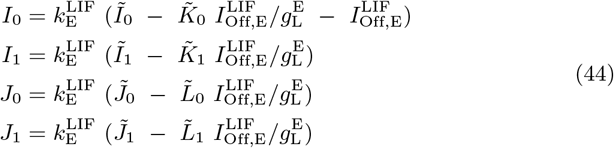

with the *I* and *S* terms on the right-hand side defined by Eqs 41 and 42.

These relationships complete the mapping to the current-based FR ring model G.

### 2.3 Simulations

We now present comparative simulations of the different models described in Section 2.1 and responding to to the same input. These simulations demonstrate the consistency of the mapping expressions, show how different behaviours of the models reveal implications of each step of the reduction, and highlight essential mechanisms for certain signatures of responses in the visual cortex.

We first set the parameters of the most complex model with excitatory and inhibitory populations distributed in 2D cortical space (model A), according to the experimental literature on single neuron properties, synaptic kinetics and connectivity (Section 4.6). The parameters of the remaining models (Section 4.6) are then calculated according to the formulas derived in Section 2.2.

#### 2.3.1 Overall transient dynamics

We compared responses of the models to a specific visual stimulus, focusing on the excitatory population firing rate. The initial conditions were the steady-state in response to a homogeneous background input of “contrast” or strength *φ*_0_. The visual stimulus of contrast *φ*_1_ is presented at *t* = 0ms with a time-dependent orientation (e.g. oriented bar, grating, edge, or more complicated scene) of 0° at *t* = 0ms, shifting to 45° at *t* = 100ms. For model A, we simulated the response of 4 identical hyper-columns (Figure 2), with an identical input for each. This stimulus provides the fundamental ”step response” of the system from the background state, followed by a step response in the orientation domain, for example following an abrupt change in the visual scene during a saccade.

We now consider the overall dynamics of the responses of the 2D model A. The spatial distributions of the excitatory population firing rate at different times are shown in Figure 6. Initially, the modeled cortical domain responds with a firing activity in the orientation columns that correspond to maximal stimulus (frames at 5-20 ms). The maximum response appears at about 20 ms; its location corrsponding to the stimulus orientation. The activity is homogeneously distributed along a radius of a hypercolumn. At later stages, the firing rate decreases (frames at 30-50 ms) with a peak located near the pinwheel center (frames at 50-90 ms). The input changes its orientation at 100 ms from 0° to 45°. The activity rapidly rotates toward the column preferring the orientation of the stimulus (frames at 105-110 ms), and even beyond (frames at 120-130 ms; see also Section 2.3.5). Later, the activity settles again in the column preferring the orientation of the stimulus with a peak near the pinwheel center (frames at 150-190 ms), similar to the activity in response to initial stimulus (compare to frames at 50-90 ms). The time-dependent profiles of the rate and voltage for the populations on a ring of radius 2/3 R centered on a pinwheel (marked as a circle in Figure 6), thus as a function of preferred orientation, are shown in Figure 7 (excitatory and inhibitory populations) and Figure 8A (excitatory population). The activity of the interneuron population is stronger and more widely distributed in orientation space than that of principal cells, as seen from comparison of Figure 7B to A, and Figure 7C to the first panel of Figure 8. This activity provides broad cortical inhibition by shunting the excitatory neurons. As a result, the excitatory activity localizes in small cortical zones as shown in frames at 50, 90, 150 and 190 ms in Figure 6, as well in the snapshots of the voltage and rate as a function of distance from the pinwheel center, shown in Figures 7D and 7E, respectively. Importantly, the steady-state location of the maximum activity in each hypercolumn corresponds to the orientation of the stimulus, and the firing is localized in 2D and orientation coordinates, i.e. the cortical response is tuned.

**Fig 6.**
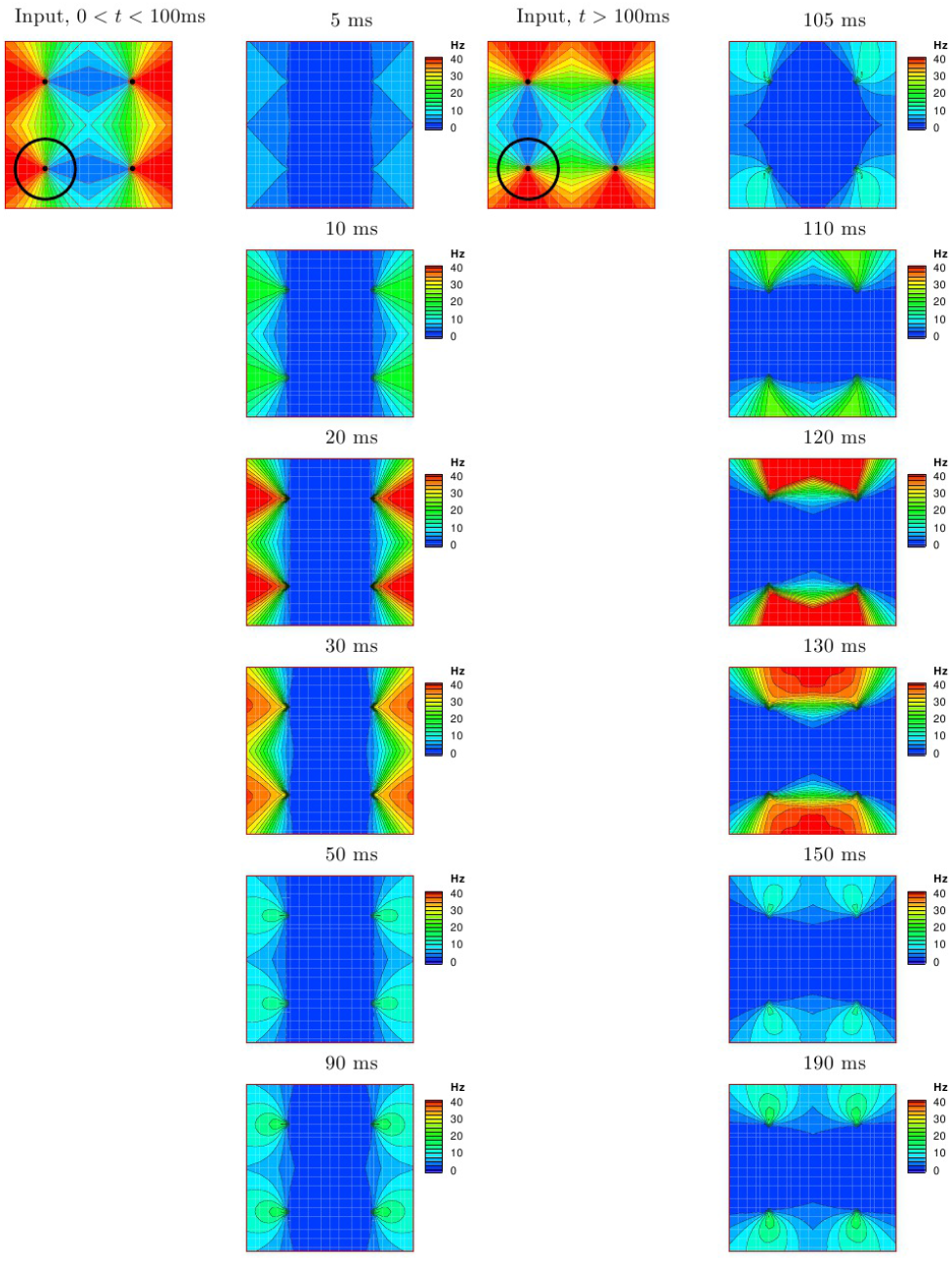
The response of model A (HH 2E EI Exp 2D) over the cortical surface (1mm^2^ containing 4 hypercolumns; the dots mark the pinwheel centers) to a 0° oriented stimulus presented at *t* = 0, and shifting to the 45° orientation at *t* = 100ms. The circle at the lower left quadrant of the input maps correspond to the activity profile of this model shown in the first panel of Figure 8.

**Fig 7.**
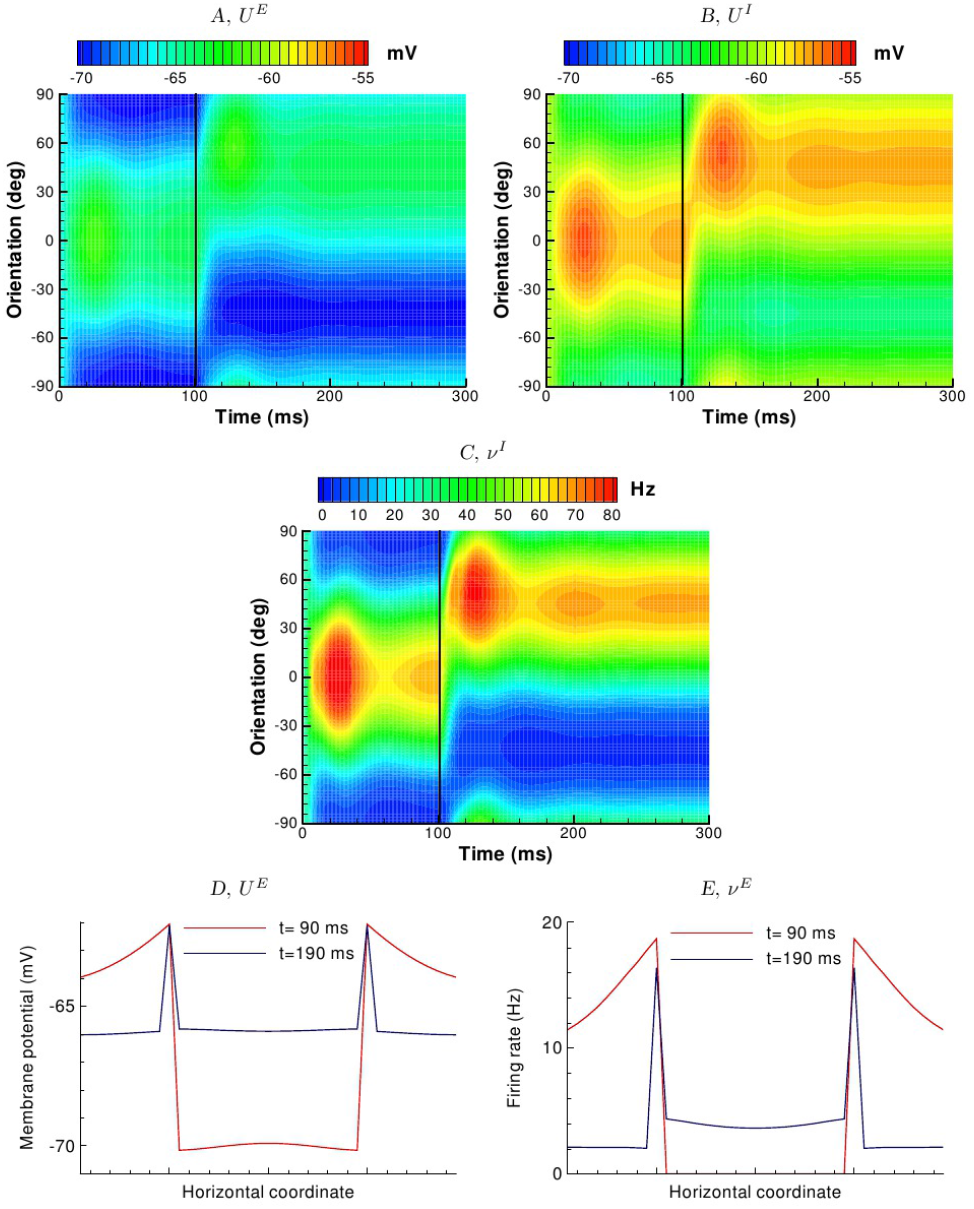
Evolution of voltage and firing rate profiles of model A (HH 2E EI Exp 2D) as a function of preferred orientation, in response to the stimulus sequence described in Figure 6. *A*, sub-threshold voltage for the excitatory population; *B*, sub-threshold voltage for the inhibitory population; *C*, inhibitory population firing rate; *D*, excitatory population membrane potential across horizontal line across the cortical surface passing through two pinwheel centers at two time moments *t* = 90 and 190ms, corresponding to the snapshots in Figure 6; *E*, excitatory population firing rate across the same line at the same time moments.

**Fig 8.**
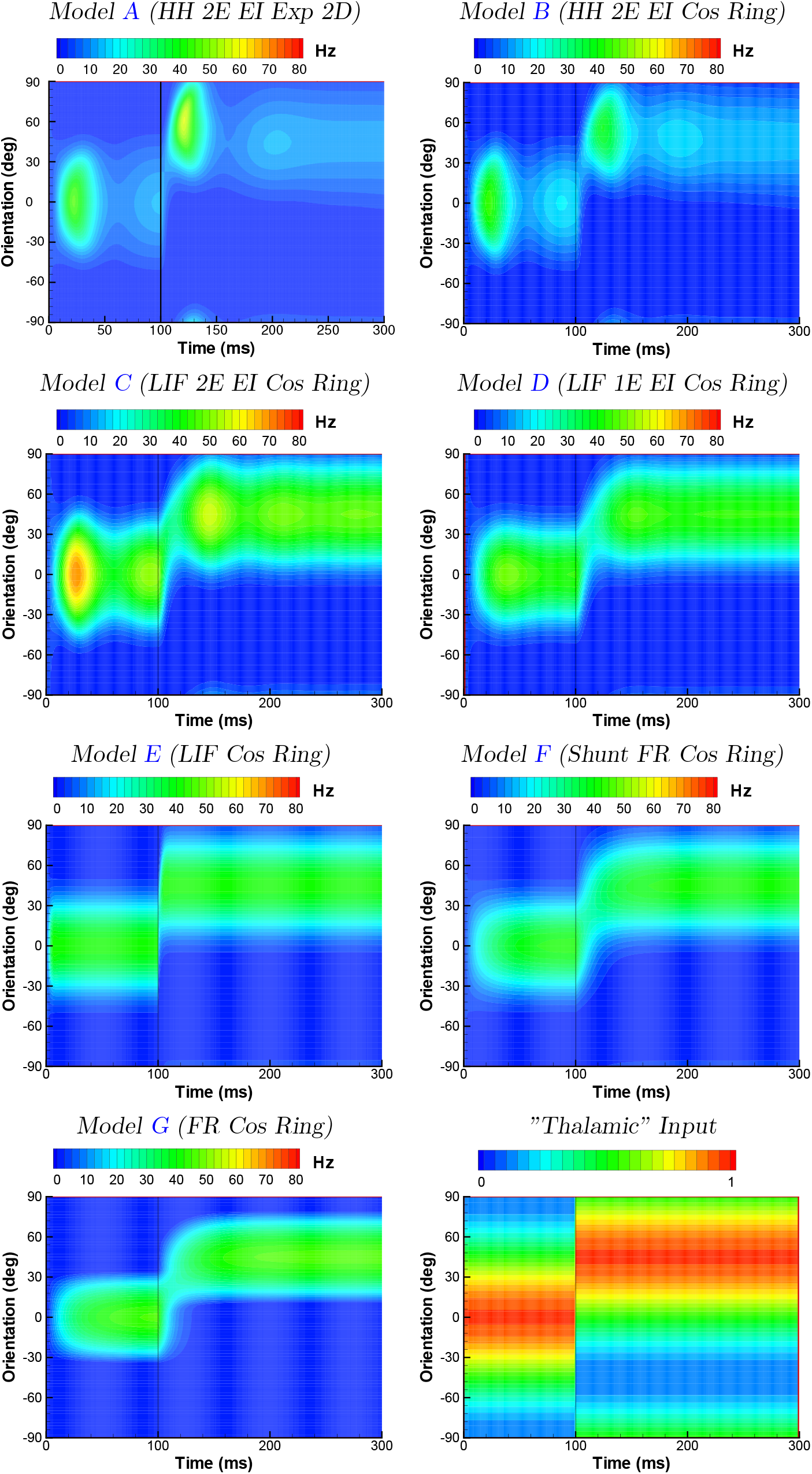
Responses of a hypercolumn as a function of preferred orientation to the stimulus sequence described in Figure 6, for the excitatory population of the various models listed in Table 1 (ref. ”Thalamic“ input panel). The activity for model A corresponds to the circle indicated in Figure 6.

The population rate as function of orientation and time for the remaining models is given in Figure 8. When considered along a ring centered on a pinwheel, the dynamics of the 2D adaptive, two population model A, is similar to the adaptive ring geometry model B, although the full dynamics of the 2D model depend on the radius of the ring, as will be discussed below. This similarity is achieved due to the mapping performed in Section 2.2.1. Both exponential and cos-shaped profiles of connection spread on a ring (Figure 3B) result in similar solutions (data not shown) seen in Figure 8 for model B. In contrast, the transient responses change significantly between the adaptive HH models A and B, and the non-adaptive LIF model C, primarily due to an overall increase in the activity in the latter model. Damped oscillations are observed to varying degrees for models A-D, being strongest for the adaptive models, and quite weak for the 1 compartment LIF EI model D. Thus, steady-state-like behaviour in the adaptive models A and B is seen only after 200ms. The non-adaptive models C and D show steady-state-like behaviour within approximately 100ms, thus observable for both stimulus orientations.

The reduction step from the 2-compartment LIF neurons of model C to the 1-compartment LIF neurons of model D, which reduces the order of the system of equations, has a more striking effect on the transient response. In particular, as noted above, oscillations following stimulus transitions are substantially reduced for the 1-compartment LIF model. A larger propensity to oscillations of the higher order model C in comparison to the lower order model D is consistent with the observation of a faster voltage response of a spatially distributed neuron to changes in synaptic current due to the electrotonic properties of the implicit cable structure. Following the change in the stimulus at 100ms, both models show a continuous shift of the response in orientation space, or a virtual rotation, as discussed below.

The subsequent model E incorporates two more simplifications by reducing the number of populations from two to one, and assuming instantaneous synaptic kinetics. These lead to quite different transient behavior, with a much more rapid establishment of the steady states, and the disappearance of oscillations and virtual rotation. Why is the response of model E much more rapid than the previous in models, even though the essential cellular time constant is similar, i.e. given by the membrane time constant which scales the rate of the voltage integration in response to a change in the input current? To explain, we note that the firing rate of model E effectively depends on both the voltage relative to threshold as well as the change in the voltage [23]. The result is a rapid response of the population to an abruptly changing stimulus [24], [25] and explains the near-instantaneous response in our simulations.

Further reduction to the FR-model with shunt, model F, with its time constant equal to the membrane time constant, leads to the delays of the responses. The reduction implies the substitution of the statistically precise consideration of neuronal states in each population by a statistically approximate approach. Namely, the Fokker-Planck-based and CBRD-based methods of models A-D (see Methods) are substituted by the FR-model that is strictly valid only at steady-states of a population. For nonstationary regimes the FR-model provides an approximation that filters the firing rate with some characteristic time constant of the filtering. By chance, given the various parameters of the models, the time constant of model F is close to the synaptic time constants, thus the transient solution is again as smooth as the solutions of models A-D. The small residual difference between the responses of the FR-ring models with and without shunt (models F and G) is explained by the approximation of the dependence of the rate on the current and conductance by the piecewise linear function (Eq 26).

#### 2.3.2 Sharpening of output tuning vs. input tuning

Our analysis continues with the steady-state response of the networks, taken as the activity profiles at *t* = 300 ms, in terms of the sharpening between input and output, and of contrast-invariance of the output tuning curves.

A fundamental aspect of the orientation selectivity in primary visual cortex is that this functional property is essentially absent in individual inputs from the thalamus. Various theories have been proposed to account for the emergence of tuning in cortical neurons, but a common assumption is that there is an initial bias in the spatial retinotopic footprint of the feedforward pathways of cortical neurons, and this bias is strengthened in terms of a sharper orientation tuning in the cortical receptive fields ([26], [27], [28]). In the context of the models described here, this bias is encapsulated by the constant plus cosine form of the thalamic input.

We consider this input-output sharpening by examining the steady-state solutions of the models (Figure 9). All the models show orientation tuning sharper than the input tuning; furthermore, the sharpening increases as the models simplify.

**Fig 9.**
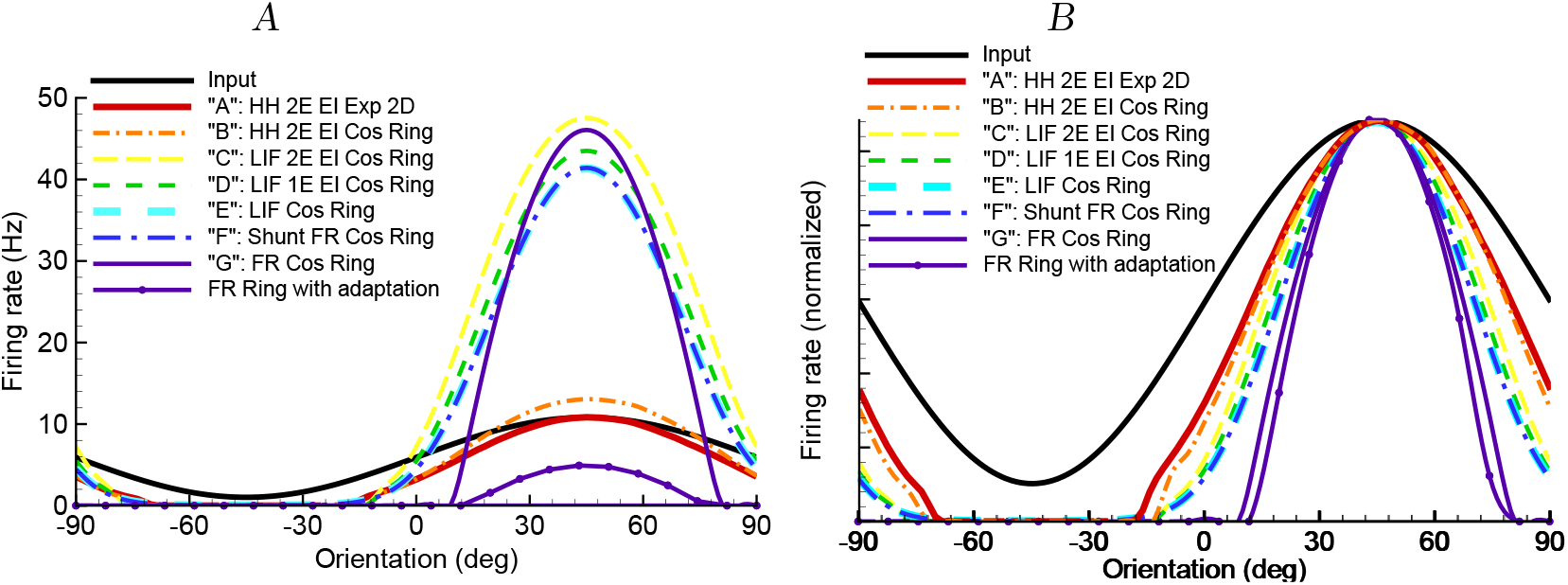
The tuning curve for the input compared to steady-state solutions for the firing rate calculated for each of the models of Table 1, obtained as cross-sections at *t* = 300ms of the plots shown in Figure 8 (un-scaled in *A*, and normalized in *B*). The profile with the smallest amplitude corresponds to the simulation by the ”adaptive” firing-rate ring model G as described in the text and indicated in Figure 10. The half-widths at half-maximums are 38° for model A, 36° for model B, 31° for model C, 28° for model D, 27° for models E and F, 24° for model G, and 22° for model G with the gain rescaled to account for adaptation.

The canonical FR ring model G has provided a semi-analytic framework for accounting for this sharpening in terms of the strength and tuning of intracortical connectivity. In this model the non-tuned inhibition is effectively stronger as *J*_0_ decreases, whereas the tuned excitation is stronger as *J*_1_ increases. As presented above, the derived values for *J*_0_ and *J*_1_ correspond to the ”marginal” domain (Figure 10) which in turn predicts enhanced output sharpening. In contrast, while the analysis described above suggests that the 2D HH model A is operating in the homogeneous domain, cell thresholds and tuned ”mexican hat” inhibitory and excitatory populations (Figures 7) still allow for input-output sharpening. The prediction that adaptation underlies a shift from the marginal to the homogoneous regime of model A during the response suggests that sharpening of the phasic response should be greater than the tonic response, as reported by measurements of orientation-tuning dynamics in the macaque [29].

**Fig 10.**
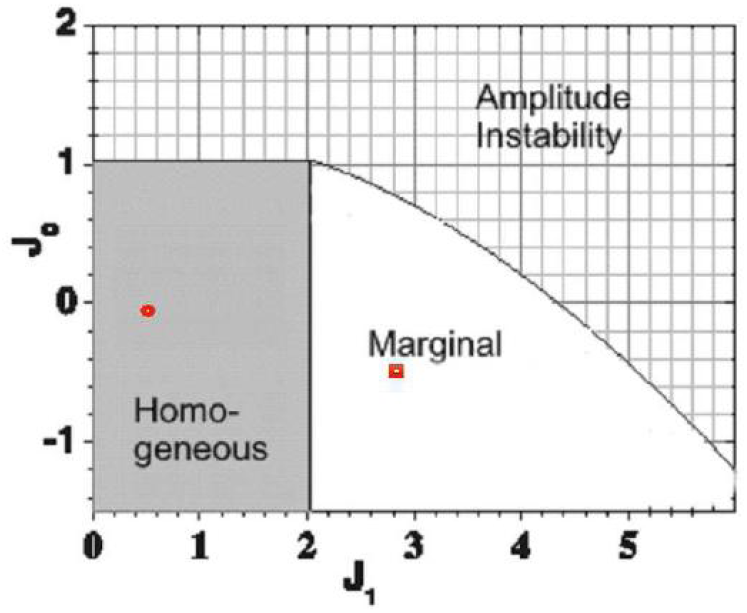
The diagram of the steady-state solutions of the canonical firing-rate ring model G on the plane of its parameters *J*_0_ and *J*_1_(adapted from [2]). Amplitude instability corresponds to the state where the activity in the ring increases without any possibility to regulate it. Homogeneous phase (“feedforward” hypothesis) corresponds to a state of weak interactions. The activity in the ring follows directly from the thalamic input, apart from a threshold non-linearity. Marginal phase (“recurrent” hypothesis) corresponds to a state where only a tuned activity profile is stable, partially but not completely determined by the input shape and dynamics. This state occurs for sufficiently strong recurrent tuned inputs (*J*_1_) and, to a lesser extent, with sufficiently strong inhibition (*J*_0_). The small square marks the parameters found by the mapping expressions for model G (*I*_0_ = −26.5, *I*_1_ = 44, *J*_0_ = −0.52, *J*_1_ = 2.8). The small circle in the homogeneous state corresponds to a variation of model G where the value of 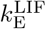 was derived from the adaptive HH excitatory cell model (*I*_0_ = −3.55, *I*_1_ = 7.4, *J*_0_ = −0.063, *J*_1_ = 0.46).

The impact of spike threshold is also seen in the simulations. When comparing subthreshold voltage and spikes, experimental investigations have reported that the tuning of the former is broader than the latter [30], [31], [32], [28], [33] et al., which can be explained simply by firing threshold. This behaviour is replicated in the the HH and LIF models which explicitly consider intracellular dynamics and spike threshold (compare for example the first panel in Figure 8, and Figure 7, for the 2D HH model A).

Figure 9 also confirms the sharp increase in the response with the removal of adaptation due to slow M- and AHP-channels, and thus the large increase in excitatory cell gain as discussed earlier (compare models A and B, with models C through G). Indeed, the stationary solutions of the non-adaptive models are quite similar, supporting the assumptions made in Section 2.2.4.

Note that the 2D and ring HH adaptive models A and B give small amplitude steady-state profiles in orientation space (Figure 9A) that are qualitatively consistent with the solution for the “adaptive” firing-rate ring model described above. However, quantitative comparison in this case, especially for output sharpening, is problematic, mainly because for the adaptive neuron the rate-current-conductance function is no longer scaled by Eq 24, and is not approximated by Eq 29.

#### 2.3.3 Contrast invariance

Contrast invariance, which is the qualitative maintenance of the output tuning sharpening with respect to the input tuning over a range of input strengths, is another classical property of neurons in primary visual cortex described by experimental and theoretical studies [34]. Attractor dynamics of the neuronal network have been proposed as the underlying mechanism for this functional property; thus the marginal domain of the canonical FR ring-model model G predicts contrast-invariance, and this property is conserved for the LIF-based ring-models E, D and C. In comparison, the fact that the adaptive version of the G model is in the homogeneous domain suggests that the adaptive models A and B will not show contrast-invariance. Figure 11 confirms these predictions for models A and D.

**Fig 11.**
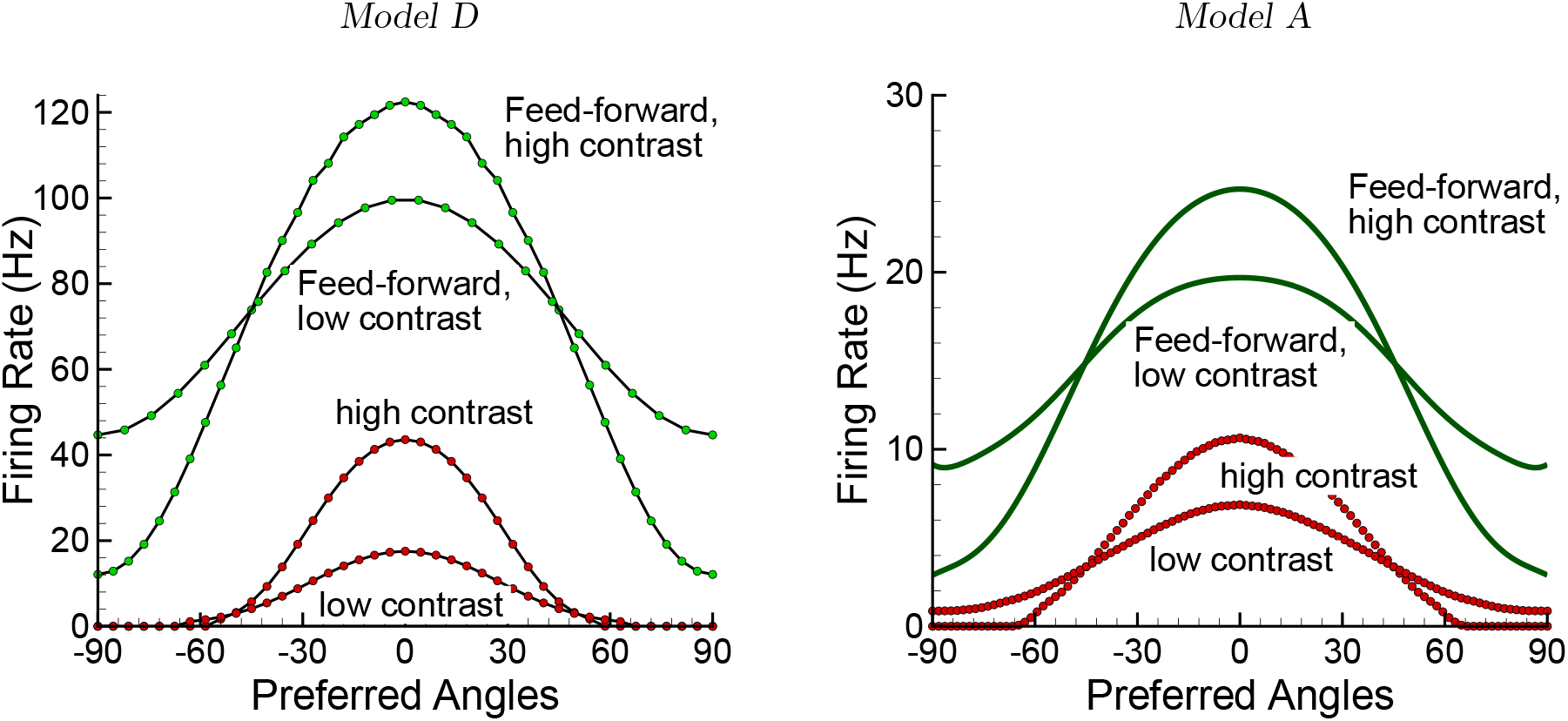
Contrast invariance for the excitatory population firing rate (red) in models D (left) and A (right), compared with the responses (green) of feed-forward versions of each model 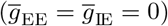 to distinguish the contribution of intracortical pathways. The “weak” and “high” contrast values were 0.4 and 0.8, respectively. As described in the text, the higher gain of model D as compared to model A, as seen in the different response magnitudes, account for the marginal phase behavior of the former, thus contrast-invariance, and the homogeneous phase behavior of the latter, thus no contrast-invariance. The intracortical connections serve to increase input-output sharpening for both models.

The models predict therefore that firing adaptation of excitatory neurons is sufficient to cause a dynamic qualitative evolution of the visual response, specifically that contrast-invariance should be observed essentially during the transient phase of visual reponses, being reduced or absent during adapted (long term) responses. Similarly for input-output sharpening, the experimental prediction is thus that the phasic response provides the basis for measured contrast-invariance; on the other hand considering the tonic response alone will show reduced or absent contrast-invariance.

#### 2.3.4 Virtual rotation

In their analysis of the canonical FR ring-model, Ben-Yishai and co-workers [1] introduced the concept of virtual rotation in the marginal phase regime, defined as the non-instantaneous movement of the peak activity following a new stimulus. They suggested that virtual rotation may be implicated in the psychophysical phenomena of mental rotation, the perception of three dimensional attributes after discrete views of an object from different angles. From the point of view of analysis, virtual rotation is one example of attractor dynamics, which in turn may provide a fundamental signature of different cortical architectures.

The stimulus sequence that we use here is particularly appropriate for examining this phenomenon, and can be appreciated in Figure 12, which traces the center of mass of the population firing rate over time. As expected given its position in the marginal domain, the canonical FR ring model G here shows virtual rotation, thus a time constant for the shift of orientation preference of approximately 30ms, as do the LIF and FR-w/shunt models D, E and F. This effect is weaker, thus the shift in orientation faster, in the non-adaptive 2 compartment LIF model C, and is essentally absent in the adaptive HH models A and B.

**Fig 12.**
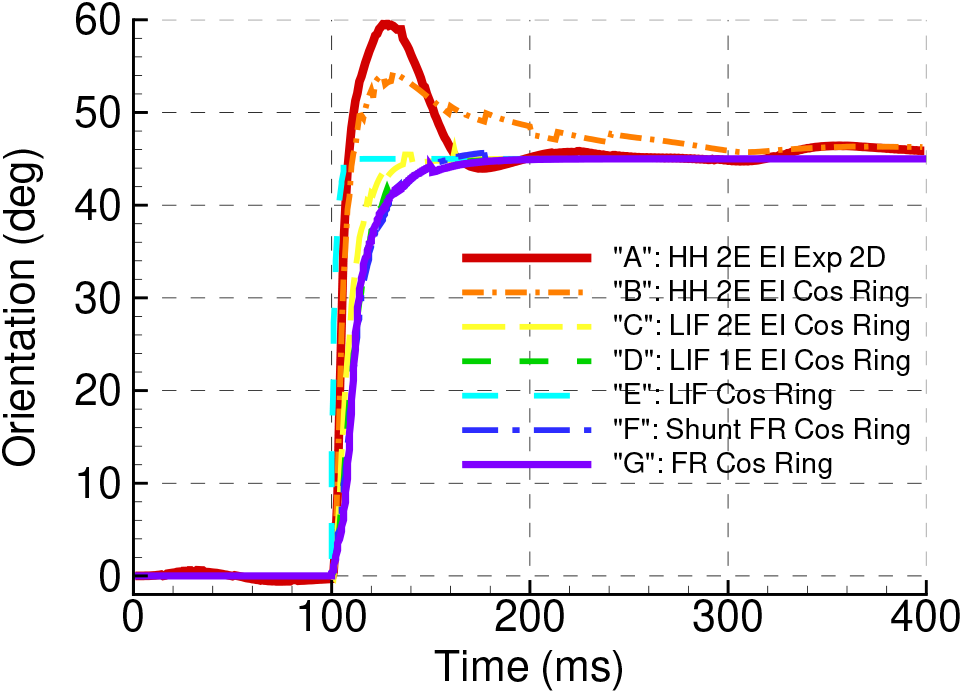
The traces of the center of mass of the population firing rate over time, calculated for all models, according to Figures 7 and 8. Note the tilt after-effect, thus an overshoot of the response tuning, and virtual rotation, thus a lag in the tuning, after the abrupt shift in stimulus orientation at 100ms.

#### 2.3.5 Tilt after effect

The tilt after effect [35], [36], [37] refers to a perceptual bias of an oriented stimulus away from the orientation of a preceding stimulus. Electrophysiological studies indicate that this effect arises, at least in part, from primary visual cortex [38], [39], and this hypothesis has been further studied with models of V1 (e.g. [40]).

As the case for virtual rotation above, the stimulus sequence examined here is well suited for studying the tilt after effect. In our models this effect is seen to varying degrees in Figure 8, as an overshoot of the maximum activity beyond the orientation of the second stimulus. Specifically, this effect is seen clearly between roughly *t* = 110 and 150 ms in the 2D and ring adaptive HH models A and B, whereas removing adaptation in the subsequent LIF ring model C suppresses the effect. The physiological studies of Dragoi and colleagues [41] have confirmed a crucial role of adaptation both for suppressing the response to the new stimulus on the side near the first stimulus, and shifting its peak activity. Inspired by these findings, the modeling study of Jin et al [40] focused in part on the impact of adaptation mechanisms on the time scale of minutes. This can be compared to the adaptation mechanisms used in the present study (M and AHP currents) which have time scales on the order of one second or less; indeed, as Jin et al note, fast adaptation (as studied by [42], [41], [43], [44]) gives a weak tilt after effect [45]. Nevertheless, the qualitative demonstration of the effect in the models examined here is consistent with additional and longer lasting adaptation mechanisms, and provide a basis for exploring this phenomena with biophysically-constrained population models.

#### 2.3.6 Spatial distribution effects

The consideration of a 2D cortical geometry can be related to two aspects of the model behaviors shown here. First, in the full 2D model A there is a gradient as a function of distance from the pinwheel centers in the steady-state or quasi-steady-state responses to both stimulus orientations, thus at *t* = 90ms and *t* = 190ms (Figures 6 and 7). This effect can be explained by the distribution of inhibition along the vector towards a pinwheel center. Thus, neurons far from the pinwheel center receive nearly circularly symmetric inhibition, whereas those closer to the center receive inhibition increasingly limited to the side away from the center. Of course, as stated this effect derives directly from the assumption of homogenous cortical connectivity in the model, and thus may be altered with a different connection scheme. Thus the model predicts higher activity near pinwheel centers if the input connections are homogeneously distributed along the radius on a scale of a single pinwheel. Second, the tilt after-effect is more pronounced for neurons remote from the pinwheel centers, as seen from curved isolines in the frames at 110, 120, 130 and 150ms. By construction, these features cannot be reproduced in any of the ring models.

In summary, the residual differences between the adaptive HH 2D model A at a fixed pinwheel radius, and it’s immediate ring version, the adaptive HH ring model B, are most likely due to the qualitative reduction of the 2D to 1D geometry as well as quantitative approximation of the synaptic projections to the ring by the cosine profile.

### 2.4 Parameter analysis of the ring models

We now present an analysis and implications of the mapped parameters of the reduced ring models, especially the canonical ring model G, in terms of the original biophysical parameters of the most detailed 2D model A.

#### 2.4.1 Reformulation of the mapping expressions

For the sake of analysis, we define several terms to simplify the expressions Eqs 41,42,44, in a manner that collates parameters of the different circuit pathways.

We first develop expressions for the 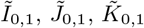 and 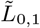 terms in Eqs 9 and 10 that define the single population conductance-based rings (models E and F, repeated here for convenience):

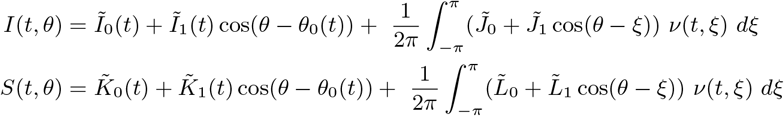

Referring to Eq 41, for the thalamic input coefficients 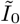 and 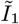, we define:

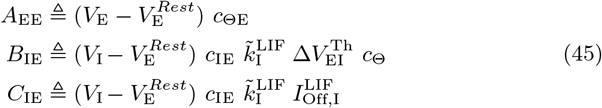

These terms characterize specific paths within the network. First, *A*_EE_ describes the explicit feedforward excitatory path from the thalamus. Recalling that 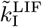 (Eq 36) reflects the effective gain of the inhibitory population (less than the actual gain of these neurons because of inhibitory-inhibitory feedback), *B*_IE_ describes the concatenation of the implicit inhibitory feedforward and recurrent connections, and *C*_IE_ characterizes the implicit inhibitory recurrent connections.

We define similar terms for the intracortical coefficients 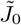 and 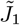:

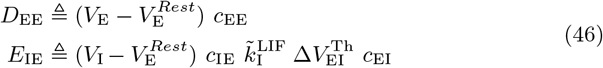

Note the similarities between *D*_EE_ and *A*_EE_, and between *E*_IE_ and *B*_IE_, respectively, in each case differing only by the final *c_ij_* term characterizing the source pathway. Thus, *D*_EE_ describes the explicit recurrent intracortical excitatory path, and *E*_IE_ describes the concatenation of the implicit inhibitory and excitatory recurrent connections of the conductance-based single population models.

Using these definitions, we now re-arrange Eqs 41:

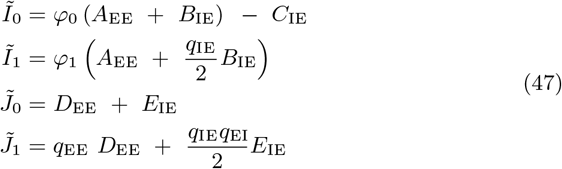

Likewise, referring to Eq 42, the corresponding conductance terms are given by:

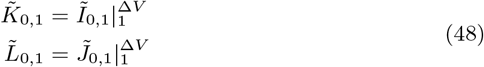

where for convenience, we now introduce the notation 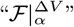 such that all the Δ*V* terms in some function 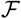 are set to *α*. For the case of Eq 48 (and later equations) we set *α* =1 (unit-less) to obtain a shorthand expression for the synaptic conductance.

We now complete the mapping by expressing the *I*_0,1_ and *J*_0,1_. terms of the current-based ring (model G) directly in terms of the biophysical parameters (Eq 3, repeated here for convenience):

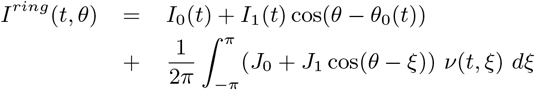

We first introduce variations on the previously defined terms (Eqs 45 and 46) characterizing specific pathways by replacing the leading 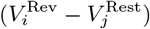 terms with corresponding 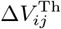 terms:

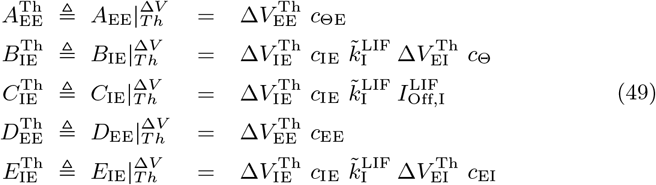

Finally, considering Eqs 44 and 47, we obtain:

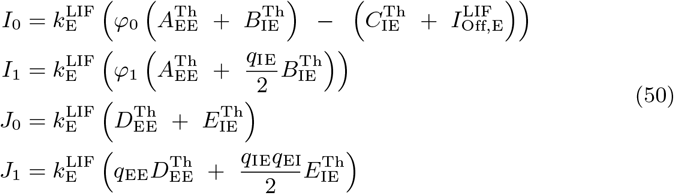

Note in particular that the pathway identification with the specific terms *A*_EE_, *B*_IE_, *C*_IE_, *D*_EE_ and *E*_IE_ of models E and F, as described in Section 2.2.4, hold precisely for the corresponding FR ring terms 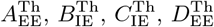 and 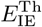 of model G.

#### 2.4.2 Feedforward inhibition

Conceptually, one can consider distinguishing functional excitatory and inhibitory synaptic pathways in terms of their thalamic, or feedforward, component, versus their intracortical component. In this context, the fact that there are no anatomical inhibitory connections from the thalamus has led to the assumption that a population of cortical inhibitory neurons, driven primarily by thalamic input, act as a surrogate for a true feedforward inhibitory pathway. The mapping to the FR ring model G allows a quantitative interpretation of this assumption.

We recall that in the original formulation of the canonical ring model, the *I*_0_ and *I*_1_ terms (Eq 3) describe the thalamic input. The mapping expressions for these terms in Eqs 50, specifically the 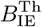 term for the FR ring, provide an explicit quantitative definition of so-called “feedforward inhibition”:

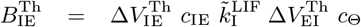

Recalling the definition of 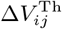 given by Eq 28, we see that 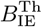 is negative because of 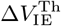. Notably, this characterizes the component of the functional thalamus-driven pathway, allied with cortically-driven inhibition (parameterized by 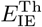), that competes against feedforward and intracortical excitation.

Finally, as seen in Eqs 50, the relative strength of the un-tuned versus tuned components of the feedforward inhibition depends on the tuning of the intracortical inhibitory-excitatory pathway, parameterized by *q*_IE_.

#### 2.4.3 Contrast and tuning of thalamic input

We now compare the thalamic input across the models, characterizing them in terms of a “contrast” parameter Ω and a tuning parameter Γ. We define Ω as the average strength of the input to the network, thus across orientation space. The response of many visual neurons is strongly related to the visual contrast of the stimulus, e.g. the response scales with the amplitude of a grating, after adaptation to its average value. The tuning parameter Γ characterizes the orientation dependence of the retino-geniculo-cortical, thus thalamic, pathway to the cortex. Indeed, as noted before, this pathway is explicitly established by the constant plus cosine form of thalamic input for all the models. For the two population models driven by thalamic firing rates, we have:

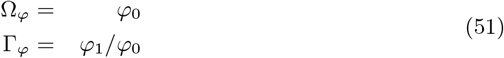

Given that firing rates must be non-negative, note that the constant plus cosine form (1 + *cos*(*θ*)) imposes the constraint 0 ≤ Γ_*φ*_ ≤ 1.

We follow the same form for defining input contrast and tuning of the single population conductance-based ring models, but now introducing separate definitions for the current and conductance inputs:

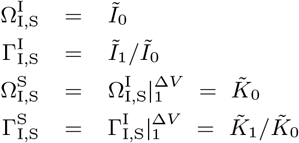

With these definitions, we now establish their relation with the contrast and tuning of the thalamic firing rate input for the two population models. Expanding 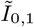 (ref. Eqs 47), and considering that *φ*_1_ = Γ_*φ*_Ω_*φ*_:

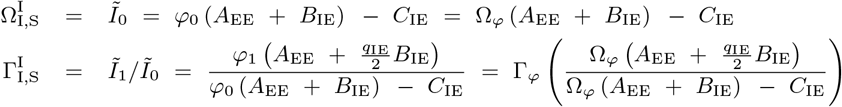

We see that there is not a one-to-one correspondance between the measures for the different architectures. First, for zero “true” thalamic input defined by firing rates, the equivalent contrast of the current and conductance inputs is − *C*_IE_ and 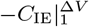, respectively, which reflect the subthreshold constant terms that result from eliminating the inhibitory population. Second, these offsets make the tunings of the current and conductance inputs dependent on the contrast of the thalamic firing rates, whereas in princple tuning is defined purely by anatomical connectivity, regardless of contrast. Finally, the coefficients for the input rates *φ*_1_ and *φ*_0_ in the tuning expressions are not equal unless 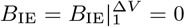 (e.g. zero thalamic input to the inhibitory population of the two population models, thus *c*_Θ_ = 0).

If we assume that inhibition is purely shunting (*V_I_* = *V*^Rest^ and therefore *B*_IE_ = *C*_IE_ = 0), then the contrast and tuning for the current input, at least, simplifies:

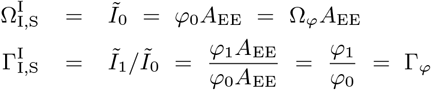

In this case, compared to the two population models A-D, we see that the “contrast” of the conductance-based single population models E and F are the same apart from a scaling term, and more importantly the tunings are identical.

We now consider contrast Ω_FR_ and tuning Γ_FR_ for the canonical FR ring model G, again following the definitions for the other models:

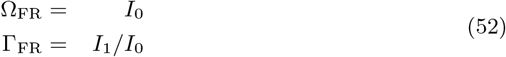

To simplify the previous expressions for *I*_0,1_ (ref. Eqs 50), we define:

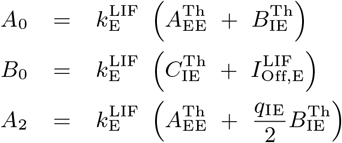

Applied to the expressions for *I*_0_ and *I*_1_, we obtain:

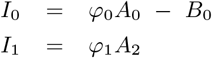

We re-express these equations, now in terms of the original definitions of contrast and tuning (Eq 51):

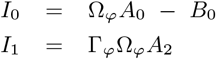

Given the definitions of contrast and tuning for the canonical FR ring (Eqs 52), we obtain:

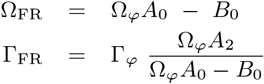

As before with the input terms of the conductance-based single population models (except for the case of pure “shunting” inhibition), the contrast of the input to the FR ring model is a non-linear function of the contrast as defined for the original two population models.

We can supply some constraints on *A*_0_ and *A*_2_ by asserting that intracortical connection weights are non-negative, thus for the constant plus cosine approximation 0 ≤ *q_i,j_* ≤ 1. This gives *A*_0_ ≤ *A*_2_, and therefore:

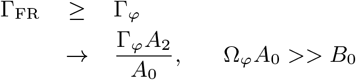

Note that Γ_FR_ diverges as Ω*_φ_* approaches *B*_0_/*A*_0_, which in turn is equal to the threshold firing input:

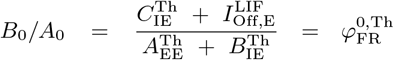

The distortion of the input contrast and tuning when compared to the thalamic firing rates is due to *B*_0_, which encapsulates the impact of the original two firing thresholds (of the HH excitatory and inhibitory neurons) on the final mapping. As one exercise, note that since 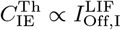, at the limit of 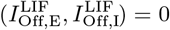, then *B*_0_ = 0.

Of particular interest are the quantitative values of the input to FR ring model G, specifically that there is a strong negative current away from the preferred orientation. This aspect runs contrary to the general assumption of the canonical FR ring model, thus that the excitatory-only thalamic pathway to the cortex constrains the (current) input to this model to be non-negative. This result arises from the effect of the cortical connections on the thalamic input terms as defined by the mapping. This is first seen in the mapping of the two population conductance models to the single population conductance models, specifically as expressed in the equations describing the cortical input, thus for 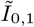 (Eqs 41) and, by extension, 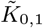. In particular, these expressions include negative terms that depend on the strength of the intracortical connections between inhibitory and excitatory populations, thus directly proportional to *c*_IE_ (for both 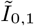 and 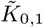) and the weighting parameter *q*_IE_ for the cosine approximation (for 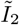 and 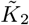). Note that these same dependencies hold for the intracortical connectivity parameters 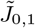 (ref. Eqs 41) and by extension 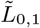. This ”crosstalk“ onto the thalamic input is conserved in the final mapping between the single population conductance models and the current-based FR ring, as seen in the equations for *I*_0,1_ (ref. Eqs 44) which depend directly on 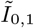 and 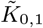. As with the single population conductance models, there are similar dependencies for the intracortical parameters *J*_0,1_ (Eqs 44), which depend directly on 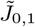 and 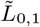.

We propose that the quantitative mapping developed here provides a novel explicit interpretation of the concept of “feedforward-inhibition”, as introduced in Section 2.4.2, thus corresponding to the negative input current shown here.

#### 2.4.4 Intra-cortical connections and the dynamical steady-state behaviour of the network

We have shown that the diagram of the steady-state solutions mapping formulas provide an interpretation of the coefficients *J*_0_ and *J*_1_ of the FR ring model in terms of quantitative biophysical, thus synaptic and neuronal, parameters. This then allows an analysis in terms of the phase-plane of the FR model, as described by [2], shown in Figure 10. From this plot we see that the parameter values of the derived ring model G place it in the marginal phase, and thus allowing for attractor dynamics.

The mapping expressions (Eqs 50) show that all the firing-rate ring model parameters, *I*_0_, *I*_1_, *J*_0_ and *J*_1_, are proportional to the gain of the excitatory neuron population, 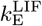. This underlines the fundamental role of 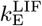, a purely local cellular, and not network, property, with respect to the phase space of the canonical FR model: a linear trajectory in this space necessarily begins in the homogeneous regime (which includes the origin). Depending on the relative values of the network components 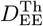 and 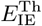 (parameterizing the recurrent excitatory and inhibitory pathways, respectively, and weighted by the *q_ij_* parameters in the case of *J*_1_), the trajectory subsequently passes either into the marginal or unstable regime and remains there, or first into the marginal and then the unstable regimes.

As constrained by the biophysical parameters, this dependence is significant across the model spectrum described here, most crucially on *J*_1_. Indeed the removal of adaptation between models B and C significantly increases the excitatory cell gain (from 0.069 Hz/pA for the adaptive neuron to 0.23 Hz/pA for non-adaptive neuron, ref. Figure 5), thus 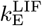. Derivation of an “adaptive” version of model G underlines this significance, where the value of 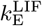 derived from the adaptive HH excitatory cell model puts the model in the homogeneous state (Figure 10). As presented later, this qualititative difference is manifested in several ways by the simulations. The effect of rate-dependent adaptation can be qualitatively compared to the effect of tuned inhibition which decreases *J*_1_, both providing negative contributions to the terms with the rate in the right hand part of Eq 1. The consequences of an increase of the adaptation or the tuned inhibition are similar, however the adaptation is more concentrated than the cosine-like tuned inhibition, and hence is more efficient.

We now examine 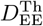 and 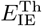 in more detail. As the case for constraints on the biophysical terms 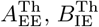 and 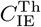 described earlier, reasonable assumptions on the reversal potentials for the excitatory and inhibitory synapses imply:

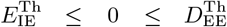

Thus, the expressions for *J*_0_ and *J*_1_ show explicitly the direct competition between the net strengths of excitatory and inhibitory recurrent pathways in determining the behavior in phase space, specifically both the quadrant that the network is restricted to (given by the signs of *J*_0_ and *J*_1_) and the sensitivity of the location in phase space as a function of the excitatory gain 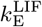. For *J*_0_ the competition between the pathways is independent of the anatomical spread of the various connections. In contrast, for *J*_1_ the contribution of the excitatory recurrent network is weighted by its characteristic anatomical tuning parameter *q*_EE_, while the inhibitory recurrent network term is weighted by the product of *q*_IE_ and *q*_EI_, reflecting the effective convolution of the recurrent excitatory-inhibitory and inhibitory-excitatory pathways.

The sign of *J*_1_ establishes the qualitative relation between the intrinsic stable attractor states and the input. Thus, a positive *J*_1_ causes the attractor steady-state to line up with the stimulus, while a negative *J*_1_ tends to cause the attractor steady-state to be orthogonal to the stimulus. Note that this applies strictly to the limit case of homogenous input, because the actual steady-state of the network is established by both the inherent attractor properties and the quantitative value of the stimulus. For illustration, we consider the limit cases for a network with flat excitatory recurrent connections (i.e. *q*_EE_ = 0), flat inhibitory recurrent connections (i.e. *q*_EI_ = 0 or *q*_IE_ =0), or both. Noting that *J*_0_ is unaffected by these parameters, for the first condition:

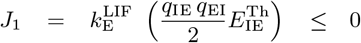

and thus the network is constrained to the left half of the phase space.

For a network with flat inhibitory recurrent connections:

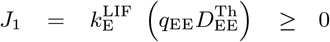

and thus the network is constrained to the right half of the phase space. These limit cases provide a more general interpretation, thus that the effective width of intracortical inhibition must be broader than that for intracortical excitation to allow the attractor properties to complement the input, and vica-versa.

Finally, if both intracortical pathways are flat, then *J*_1_ = 0, and the marginal regime is inaccessible. To our knowledge the possible relevance of a negative *J*_1_ to actual biological circuits has not been explored.

In the present case, the biophysical parameters establish that *J*_0_ < 0 and *J*_1_ > 0, thus limiting the network to the fourth quadrant of the phase space as a function of 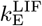. The specific mapping of the canonical FR ring parameters for the intracortical connections predict that the network is operating in the marginal phase, which is of particular interest in the context of attractor networks. More specifically, this has been used to interpret the contrast invariance of the network. However, the distortion of the input parameters of the canonical model described previously, with respect to the original thalamic input parameters *φ*_1_ and *φ*_1_, complicate the interpretation. As described earlier, a simple change of contrast Ω_*φ*_ = *φ*_0_ for the full model while maintaining the same tuning (defined as Γ_*φ*_. = *φ*_1_/*φ*_0_), maps to an input for the canonical FR ring model with a different tuning.

## 3 Discussion

### 3.1 Mapping assumptions

We can speculate on the implications of the several steps to eliminate specific non-linearities in order to achieve the final linear model (apart from the FR threshold). First, the role of inhibition may be under-estimated for two reasons. The experimental evidence on the impact of synaptic shunting on neuronal transfer functions is mixed, with some studies reporting a pure subtractive effect [46], while others showing a mixture of a threshold shift and gain reduction [47], [48]. The linear mapping here necessitates neglecting any gain change, implying that the gain of the current-based FR ring may be over-estimated, particularly in the face of strong recurrent inhibition. Furthermore, the anatomical spread of inhibitory-inhibitory connections in cortex is non-negligible on the scale of the hyper-column. The mapping requires that this spread be ignored to allow the elimination of the explicit inhibitory population, again with the result that the impact of inhibition in the final model may be under-estimated.

Second, the role of excitation within cortex may be overestimated for the following reason. A necessary assumption here is that the inhibitory population is always suprathreshold, allowing the actual threshold of this population to be ignored. However, given the large dynamic range of cortical dynamics, it is likely that different populations of neurons will be sub-threshold at different moments, especially for relatively small, but realistic, stimuli. In the context of the single population models, this implies that with small stimuli these models will transform inhibitory paths to excitatory (e.g. at the tails of the activity profile), effectively equivalent to a broader anatomical tuning of the excitatory recurrent connections.

Finally, the post-synaptic response in biological neurons to an incoming spike train saturates for several reasons, most importantly because of the reduction in driving force as the post-synaptic potential approaches the reversal potential, the finite number of post-synaptic receptor/channel complexes, not to mention mechanisms such as synaptic adaptation [49]. Indeed, only the first mechanism is considered in the two population models (in part in the single population LIF model for the excitatory population). The net result for the final FR ring is that the activity in the ring may be overestimated, particularly the peak at the preferred orientation.

### 3.2 Interpretations from the mapping

The proposed mapping allows a step-by-step fitting of parameters of increasingly abstract models to experimental data. At the most abstract level, the mappings allows the interpretation of the canonical firing-rate ring model in a slightly uncommon but more rigorous way. We see, for example, that although *I*_0_ and *I*_1_ are defined as thalamic input, when mapped from the full model, both parameters depend on the inhibitory-to-excitatory intracortical connections 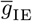 (Eqs 50, 49, 66 and 28), and therefore can not be distinguished as pure background and tuned inputs. On the other hand, study of information processing in the visual cortex by the means of the detailed model, also requires fitting of the canonical firing-rate ring model to the experimental data, in order to limit the domain of physiologically meaningful parameters’ values. This work exceeds the frame of the present paper and will be done in the future.

### 3.3 Transient behaviour of the models

After the parameters of the full model A are established, the mapping procedure provides parameters of the progressively more abstract models that are explicitly constrained by replicating the steady-state regime. At the same time, we note that the dynamics of the full model A show a rich variety of the solutions, including oscillations, waves etc. not seen in the more abstract models. Nevertheless, the systematic mappings provide constrained reference points in the parameter spaces of the models, which give essentially similar behaviors between nearest neighbors among the model hierarchy.

Generally, models with more complex dynamics possess a larger variety of solutions, like the FR ring model with a delay [5] in comparison to the canonical ring model [2]. We note that some predictions of the marginal domain of the canonical FR ring-model are maintained throughout the model hierarchy, specifically sharpening of the output tuning, while others fail for more complex models, specifically contrast invariance and virtual rotation. Summarizing the most significant differences between the models, we underline that the reduced models do not reproduce the ripples of activity provided by the synchronization of spikes and refractory effect after stimulus presentation, nor the effects of firing adaptation. We failed to observe any significant delay in the detailed models due to virtual rotation effect predicted by the reduced models, instead the reaction was fast. For the parameter sets corresponding to more prominent virtual rotation in the canonical FR-ring model we observed oscillations in the detailed models (data not shown). These observations warn that an oversimplification may lead to misleading conclusions in regards to transient behaviour of the reduced models.

### 3.4 Conclusions

We conclude that the consistency of the activity patterns of the considered models support the validity of the obtained mapping expressions, and thus permits understanding the significance of the various assumptions made during the derivation of the models’ equations. The constructed hierarchy of models can therefore serve as a useful instrument for the fitting of mathematical models [50], [51], [52] to experimental data and their subsequent analysis. In particular, the architecture of the visual cortex ring model has been recapitulated in models of other systems, either generalized [53] or specific, for example head direction cells [54], [55] and prefrontal working memory [56] in rodents, as well as navigation circuits in the fly [57], [58]. The methods developed here should be amenable towards constructing biophysically constrained abstract models of these systems.

## 4 Methods

To build a model of interacting populations of neurons we explore the probability density approach [59], [60], [11]. It is based on the equivalence of the consideration of a stochastic differential equation for a single neuron to the probabilistic consideration of neuronal density evolution in the phase space of neuronal state variables. In this framewok, simulations of the stochastic model is equivalent to consideration of a statistical ensemble of similar neurons receiving similar input signals.

### 4.1 Kolmogorov-Fokker-Planck (KFP) approach for LIF neurons with instantaneous synapses

Here we review the probabilistic evaluation of an infinite population of 1-compartment LIF neurons with Gaussian membrane current noise, used to obtain the simulations of the single population LIF ring model (model E) [60]. Thus, Eq 4 can be considered as a Langevein equation for a single neuron. This equation is equivalent to the KFP equation written for the probability density function, or neuronal density, in the voltage phase-space, *ρ*(*t, V*) (the temporal dependencies of the synaptic current and conductance input, *I* and *S*, are implicit):

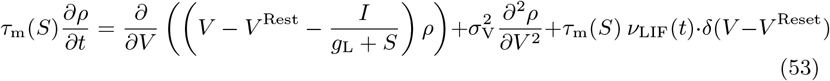

with the effective membrane time constant of the LIF neuron, *τ*_m_(*S*), given earlier by Eq 7, and the average firing rate, *ν*_LIF_(*t*), given by:

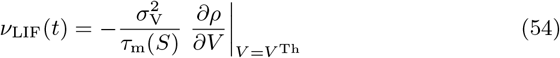

The set of Eqs 53, 54 is equivalent to an infinite set of Eqs 4. We compare the current step response of different models (Figure 13), in particular against simulations of individual LIF type neurons using a Monte-Carlo framework which provides a ”gold-standard”. The current-based FR model only roughly reproduces the transient dynamics of the Monte-Carlo simulations of LIF neurons. The KFP indeed shows convergence to the Monte-Carlo simulation of individual LIF neurons.

**Fig 13.**
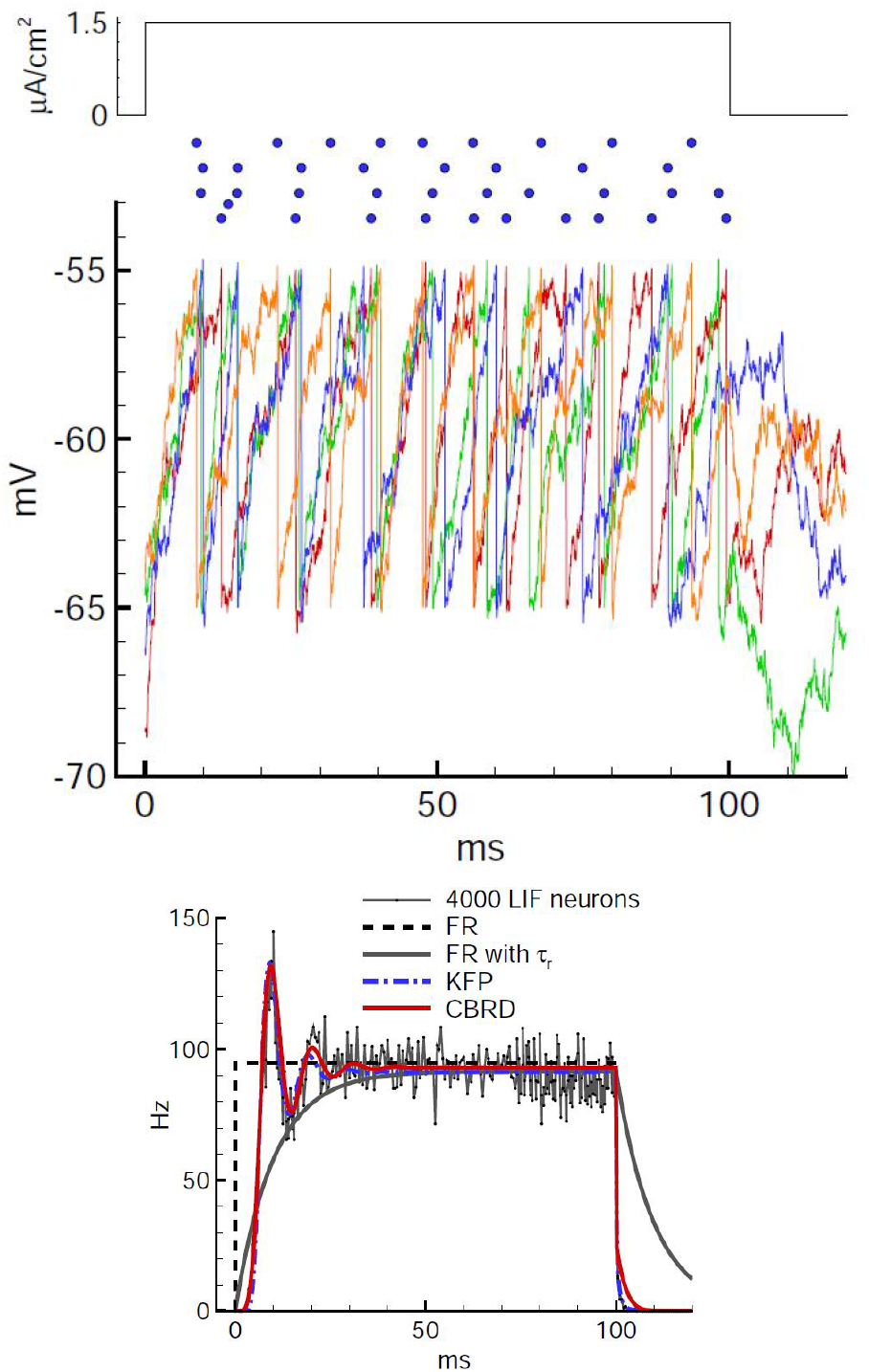
The firing of a population of noisy LIF neurons in response to current step stimulation. From top to bottom: Current step. Spike raster plot for for 4 neurons. Voltage traces for the 4 neurons. Population firing rate calculated by five models: Monte-Carlo simulation of 4000 individual trials of a noisy LIF neuron, the firing-rate (FR) model based on Eq 2, the firing-rate (FR) model with *τ_FR_* based on Eq 1, an infinite population of noisy LIF neurons evaluated with the KFP approach (Eqs 53,54) and the conductance-based refractory density (CBRD) approach (Eqs 55,57). The response of the FR with shunt cell model (not shown) is identical to the current-based FR cell model, since in both cases the transient response is fixed by *τ_FR_*, and the threshold-linear transfer function of the latter was fit to the steady-state firing rate of the LIF model, that in turn defines the former.

### 4.2 Conductance-based refractory density (CBRD) approach for multiple state variable neurons

Because of their multiple state variables, evaluating noisy neural populations with non-instantaneous synapses, multiple compartments and/or active HH channels with a KFP-based method, as described above, cannot be evaluated analytically, and thus a KFP (or Monte-Carlo) approach requires significant computational resources [61], [11]. To address this problem, we employ a conductance-based refractory density (CBRD) approach that can evaluate a broad class of models in a one dimensional phase space parameterized by the refractory time, or time since the last spike, *t*\* [17], [62], [25]. In the present case, this approach is used to evaluate all the two population models, including the HH models A and B, and the LIF models C and D.

In conjunction with a given neuron model, the CBRD approach considers a hazard function *H* which defines the probability density of firing, and describes the dynamics of a population as a whole by the probability density of neurons distributed according to their refractory times, *ρ*(*t, t*\*). In particular, the CBRD model substitutes the phenomenologically derived hazard function developed in the original refractory density approach [60] by its rigorously derived expression [17], [62]. The population model for the neuron model is expressed by a set of transport equations, one for each state variable, in partial derivatives with *t*\* and time *t* as independent variables, and thus the number of equations is directly proportional to the complexity of the model.

In order to obtain an expression for *H* in a probabilistic framework, e.g. in the presence of noise, the CBRD approach requires that the state variables be well described at any given *t*\* by a linear model around their mean value. This constraint holds for most of state variables of the HH model given their voltage-dependence and a typical dispersion of the membrane voltage of several millivolts. For the LIF model, this constraint is automatically satisfied.

In contrast, the strong nonlinear properties of sodium channels of the HH model near threshold do not meet this criterion. To address this problem, and as mentioned in Section 2.1.3, we use a threshold HH cell model without sodium channels, with spikes defined by an explicit dynamic threshold and renewal function. This model is based on three important assumptions: first, that the probability of firing can be well predicted by the dynamics of a neuron without an explicit sodium current, that the influence of strongly non-linear channel dynamics, namely the sodium current, is only significant during the spike, and third, that there is a fixed (i.e. renewal process) impact of each spike on the state variables. The last assumption allows imposed conditions on the state variables following a spike, for example reset values for the voltage and fast potassium channel gating variables (e.g. for *I*_DR_), and increments for slow potassium channel gating variables (for *I*_AHP_ and *I*_M_), as appropriate. We have shown previously that these approximations are valid in a range of neuron models [17]. The associated threshold model compares quite well to the full conductance model with respect to interspike potentials and spike time moments.

The dynamics of a given population in the CBRD model are thus described by a set of equations of the form *dY*(*t, t*\*)/*dt* = *F*(), where *Y*() includes the distribution *ρ*(*t, t*\*) of neurons with a given value of *t*\*, the average soma voltage *U*(*t, t*\*), the average dendrite voltage *U*_D_(*t, t*\*) for two compartment models, and finally the average gating particle states *x*(*t, t*\*) for HH cell models. Since the refractory time *t*\* between spikes is proportional to *t*, the total time derivative *dY*(*t*, *t*\*)/*dt* may be replaced by a sum of partial derivatives in *t* and *t*\*, i.e. *d*/*dt* = *∂*/*∂t* + *∂*/*∂t*\*.

The source term in the right-hand side of the equation for *ρ*(*t, t*\*) is the fraction of neurons crossing the threshold, given by *ρ*(*t, t*\*), multiplied by the hazard function *H*:

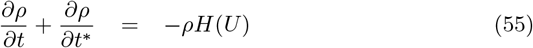

The remaining terms consist of the model details. The expressions corresponding to the cell voltages derive from those presented in Section 2.1.3, specifically the sets of Eqs 13 and 14, with the following replacements of the average voltages *U, U*_D_ and *ξ*, for *V, V*_D_ and 0, respectively:

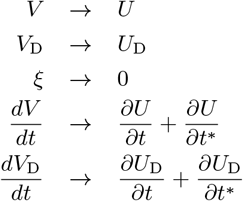

The boundary conditions for *ρ* (which also defines the population firing rate, *ν*(*t*)) and the model voltages are:

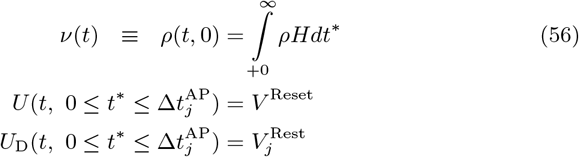

where 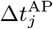 is the duration of the action potential for cell type *j* (= 0 for the LIF model; for the HH cell models see Sections 4.3 and 4.4); likewise, the somatic reset voltage 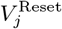 is defined also for each cell type.

The CBRD expressions for the gating particle states *x*(*t, t*\*) of the HH channel models, including their boundary conditions, are presented in the next sections (Sections 4.3 and 4.4), which in turn reference the average somatic voltage *U*.

The dynamics of the entire neural population are found by the integration of Eqs 13 and 14, with the replacements indicated above for the average voltages *U* and *U*_D_, which then defines the distribution of cell voltages across *t*\* at each time *t* (for the HH models similar equations are solved for the gating particle state variables). The effect of threshold crossing in response to the input and noise is then taken into account by the *H* function, with the integration of Eq 55 giving the distribution of *ρ* across *t*\* as well as the firing rate *ν* at time *t*.

As the source term in the density equation, the *H* function is the solution of the classical first-time threshold crossing problem for arbitrary history of stimulation. The *H* function has been semi-analytically derived from the KFP equation for voltage fluctuations, based on the following assumptions:

i. Away from threshold, voltage fluctuations due to individual noise realizations can be described by a linear equation given the mean voltage *U* and the mean membrane conductance, which are in turn given by the associated transport equations and the synaptic input. Furthermore, the evolving probability density distribution of voltage fluctuations about *U* can be described by a KFP equation.
ii. The flux across threshold described by *H* are due to two additive underlying processes: diffusion along the voltage axis due to noise, and transfer in response to the input.
iii. Diffusion due to noise dominates the firing when *U* is constant, e.g. when the input is constant. In this case the problem is described by the KFP equation with a leak and a constant threshold, denoted here as *A*. For white noise the governing equation is reduced to an ordinary differential equation with an analytical solution for the spike generation probability expressed in Kummer functions [17]. In the case of correlated (colored) noise, *A* is obtained numerically and tabulated for a range of values for *U* and the time constant of the noise correlations *τ* [62].
iv. Transfer based firing dominates when excitation starts abruptly, i.e. *U* increases with infinite rate. The initial condition is a stationary Gaussian distribution of neurons in terms of their potential *V*(*t*) − *U*(*t*). This implies that the state of zero distribution immediately following the previous spike is forgotten at the time of the next threshold crossing. As *U*(*t*) increases, the moving boundary at the threshold of fluctuations, *V*^Th^ − *U*(*t*), crosses the distribution. The flux through the boundary determines the probability of spike generation, which we denote in this case as *B*, and is obtained algebraically.

In our previous work we formulated an approximation for *H* for white Gaussian noise, as a function of the time-varying quantities that characterize the cell, including *U* (and its derivative with respect to time *t* at a given *t*\*), *σ*_V_, *V*^Th^ and the effective membrane time constant *τ*_m_ = *C*/*g*_tot_, where *g*_tot_ is the total membrane conductance:

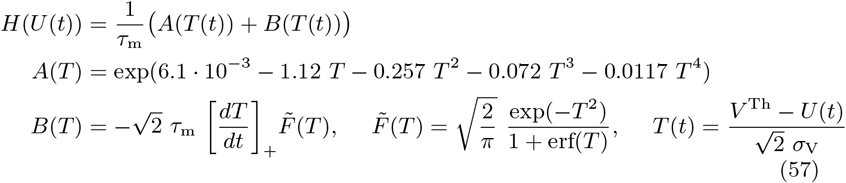

Note that the hazard function above has no free parameters, because they reflect the approximation of the first time passage solution. The dynamics of a neural continuum are thus evaluated at each time step by solving the set of one dimensional partial differential equations in terms of the refractory time, including the probability density of neurons, and the population averages of the membrane potential and channel gating variables. The CBRD model well approximates the firing rate of an infinite set of biophysically detailed neurons for an arbitrary stimulus, e.g. oscillatory input, and recurrent connections, when compared with Monte-Carlo simulations of individual neurons (see [17] and [62]). Figure 13 compares the response of a population of noisy LIF neurons, evaluated by Monte-Carlo simulations, a KFP approach (Section 4.1), and the CBRD method, with the response of the FR model with and without a time constant. Figure 14 compares the steady-state f/I characteristics as a function of synaptic conductance for a population of LIF neurons, with and without noise (Eq 5), with the result from the CBRD evaluation.

**Fig 14.**
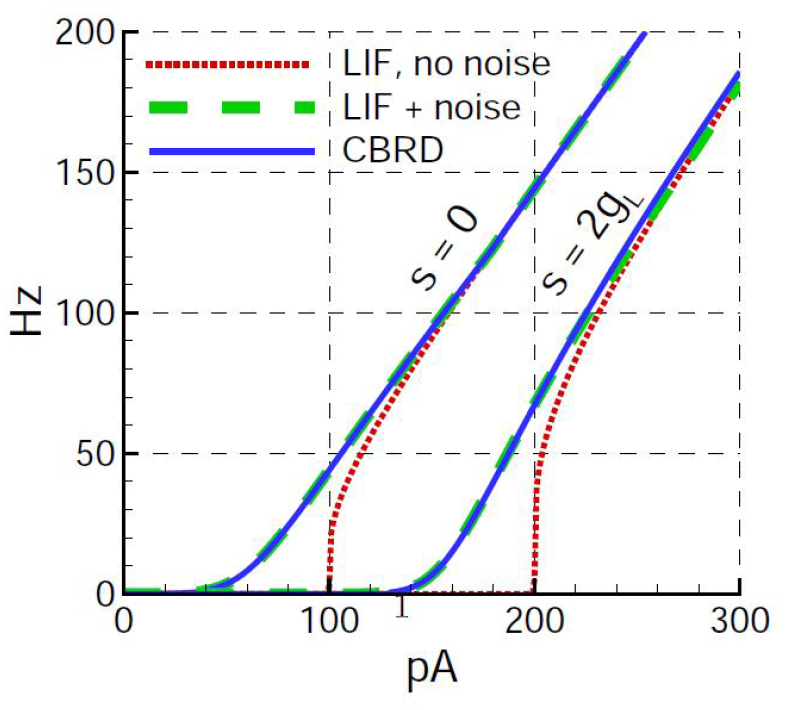
The steady-state firing rate of a population of LIF neurons as a function of the strength of the input current stimulation, with synaptic conductance *S* set to 0 and to 2*g_L_*, with and without noise (*C* =0.lnF, *g*_L_ = 10nS, *V*^Th^ − *V*^Rest^ = 10mV, *V*^Reset^ = *V*^Rest^, *τ_m_* = 10ms, *σ*_I_ = 28pA), according to Eq 5, compared with the result from the CBRD evaluation with noise.

### 4.3 Excitatory two compartment HH neuron model

Here we complete the description of the excitatory neuron model as described by Eq 14, with the average voltage *U* substituting for *V*. As stated earlier, this model incorporates several HH membrane currents, including the voltage-dependent potassium currents responsible for spike repolarization, *I*_DR_ and *I*_A_, the voltage-dependent potassium current that contributes to spike frequency adaptation, *I*_M_, the voltage-dependent cation current, *I*_H_, and calcium-dependent potassium current that also contributes to spike frequency adaptation, *I*_AHP_. The approximating formulas for the currents *I*_DR_, *I*_A_, *I*_M_ and *I*_H_ were adapted from [63]; the approximation for *I*_AHP_ was taken from [64]. The descriptions of each current are as follows.

The voltage-dependent potassium current *I*_DR_:

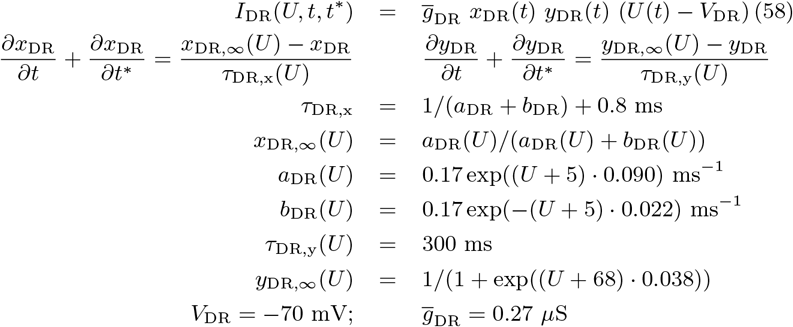

The voltage-dependent potassium current *I*_A_:

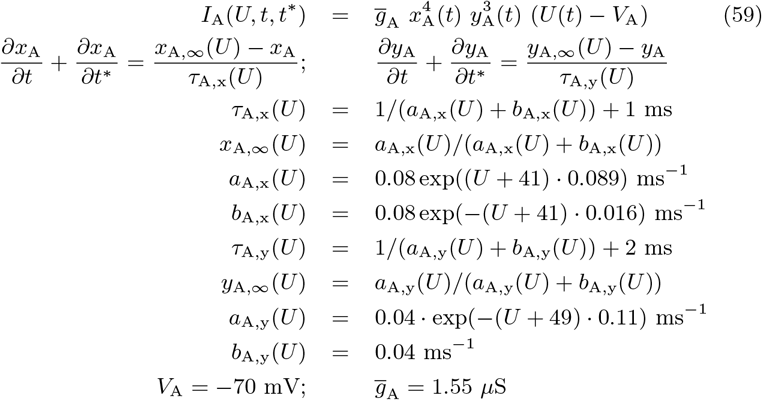

The voltage-dependent potassium current *I*_M_:

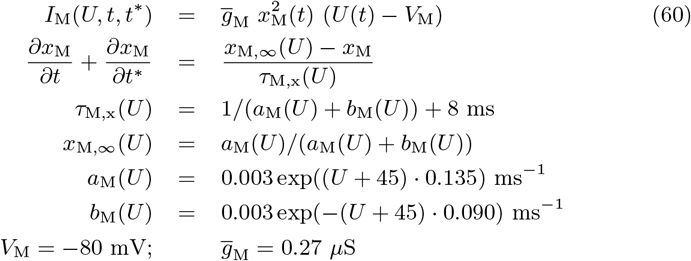

The cation current *I*_H_:

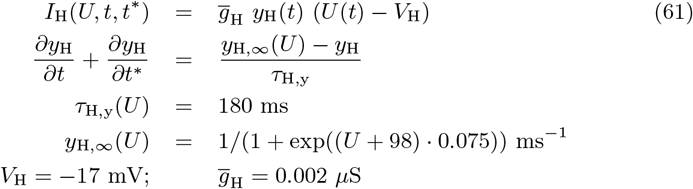

The adaptation current *I*_AHP_:

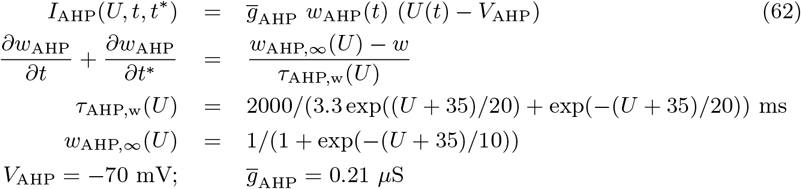

The somatic and dendritic input conductances of the two compartment model (with *L* =1), *G*_in_ and *G*_in,d_, respectively, are:

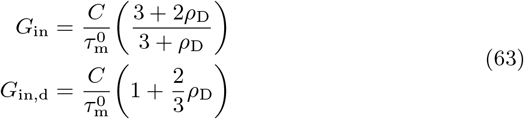

In comparison, the input conductance for a 1-compartment neuron is simply 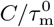. The resting time constant 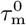 is given by:

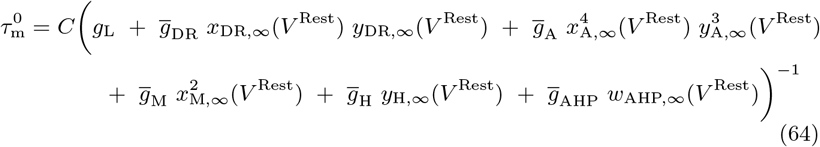

The values of the cell passive parameters were

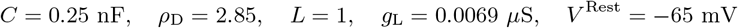

The neuron parameters provide 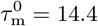 ms. The somatic input conductance *G*_in_ = 26 nS corresponds to the input resistance *R*_in_ = 39 MOhm as in [63].

According to the threshold model from [17], the duration of a spike is taken into account by introducing the time interval 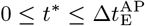 when the voltage and gating variables are reset and then kept constant, thus defining the boundary conditions of the gating particles of *I*_DR_, *I*_A_ and *I*_H_ as follows:

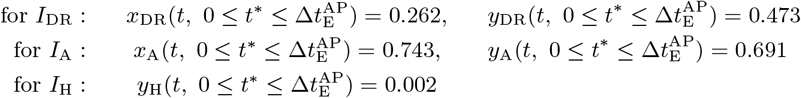

The reset values for the adaptation currents *I*_M_ and *I*_AHP_ are incremented according to their values at the peak of spike-release distribution in the *t*\*-space:

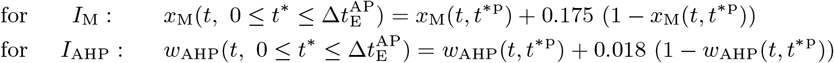

where *t*\*^*p*^ is the value of *t*\* corresponding to the maximum value of *ρ*(*t, t*\*) *H*(*t, t*\*), for *t*\* > 0.

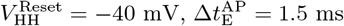. The spike threshold reference voltage in the hazard function (Eq 57) was *V*^Th^ = −57 mV, with a noise amplitude corresponding to the stationary voltage dispersion *σ*_V_ = 3 mV.

### 4.4 Inhibitory single compartment HH neuron model

Here we complete the description of the inhibitory interneuron model, according to [65], as described by Eq 13, with the average voltage *U* substituting for *V*. The membrane current for this model includes a voltage-dependent potassium current *I*_K_ according to [66]:

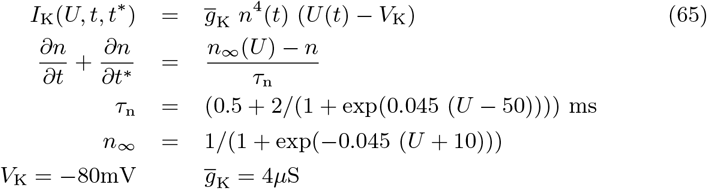

The parameters of the model are *C* = 0.1nF, *g*_L_ = 0.01*μ*S, with the boundary condition for n during the spike 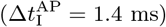 given by:

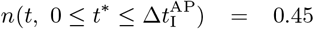

The spike threshold reference voltage, *V*^Th^, the reset voltage, 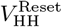, and the voltage noise parameters, *σ*_V_, in the hazard function (Eq 57) are identical to those for the excitatory cell model, above.

### 4.5 Parameterization of the second order synapse model

As described in Section 2.1.3, we include a scaling term, “synaptic capacity”, *c_ij_*, in the differential equation relating synaptic conductance and pre-synaptic firing rate, Eq 15:

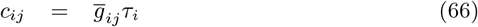

where

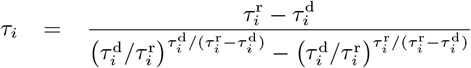

Here 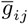 is the maximum conductance for synapse type *i* (for *i* = Θ, E, I) onto cell type *j* (for *j* = E, I), 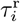 and 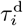 are the time constants for synapse type *i*. The time scaling factor *τ_i_* allows the maximum of *g_ij_* (*t*) to be independent of 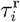 and 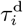 when evoked by a short pulse of *φ_ij_* (*t*) in Eq 15.

### 4.6 Model parameters

#### 4.6.1 Model A (HH 2E EI Exp 2D)

Single neuron parameters are given in Sections 4.3 and 4.4. The other parameters were set as follows:

- Synaptic parameters: 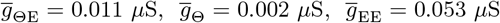, 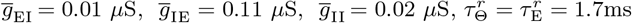, 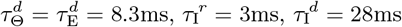, *V*_E_ = 0, *V*_I_ = −75mV.
- Spatial connections: *d*_EE_ = 100*μ*m, *d*_EI_ = 200*μ*m, *d*_IE_ = 200*μ*m, *d*_II_ = 200*μ*m.
- Region geometry: cortex region is 1mm×1mm, *R* = 250*μ*m.
- Stimulation: *φ*_0_ = 116Hz, *φ*_1_ = 96Hz, *θ*_0_ = 0° till *t* = 100ms, then *θ* = 45°.
- Numerical parameters: time step 0.1ms, spatial grid 40×40, the *t*\*-space was limited to 100ms and discretized by 100 points. The absolute refractory period was introduced by setting the term *A*(*T*) of the hazard function to zero for *t*\* < 6ms.

#### 4.6.2 Ring Models B (HH EI Cos Ring), C (LIF 2E EI Cos Ring), D (LIF 1E EI Cos Ring), E (LIF Cos Ring)

- Spatial connections: Given the value of *R* and the values of *d_ij_* for each synapse type above, fits to *w*_0_ (Eq 22) for the “exp”-profile gave 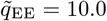, *α*^EE^ = 0.98, 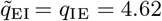, *α*^EI^ = *α*^IE^ = 1.24, while *α*^II^ = 0, 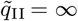 (or *φ*_II_(*t*, *θ*) = *ν*_I_(*t*, *θ*)) were set according to the assumption made in Section 2.2.4 about concentrated inhibitory re-connections. Analogously, fits to a “cos”-profile gave *q*_EE_ = 1 and *q*_EI_ = *q*_IE_ = 0.57, and *q*_II_ = ∞ (or *φ*_II_(*t*, *θ*) = *ν*_I_(*t*, *θ*)).
- Numerical parameters: ring was discretized by 40 points.

#### 4.6.3 LIF Models C (LIF 2E EI Cos Ring), D (LIF 1E EI Cos Ring), E (LIF Cos Ring)

- Excitatory model: The parameters of 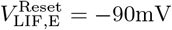, *C*_E_ = 0.27 nF are determined in Section 2.2.2, 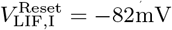 was obtain by fitting as demonstrated by Figure 4.
- For 1-compartmental neuron based model the synaptic parameters were multiplied by the factors *G*_in,d_/*G*_in_ calculated with the Eq 63 and given in the caption to Figure 4 for given value of *ρ*_D_, that gave: 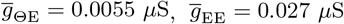.

#### 4.6.4 Models E (LIF Cos Ring) and F(Shunt FR Cos Ring)

Parameters for threshold-linear f/I approximation of 2-dimensional LIF steady-state curve (ref. Figure 5):

- Excitatory cell model: 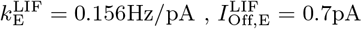
- Inhibitory cell model: 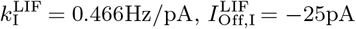

#### 4.6.5 Model E (LIF Cos Ring)

- Connections: 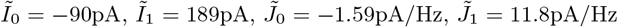, 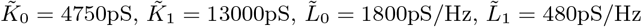 were calculated by Eqs 41,42.
- Numerical parameters: ring was discretized by 40 points, voltage space ranged from −105 (to be more negative than any expected polarization in neurons) to *V*^Th^ = −57mV was discretized by 250 points; time step 0.02ms.

#### 4.6.6 Model G (FR Cos Ring)

- The characteristic time constant of the FR model was assumed to be equal to the membrane time constant of the excitatory neurons, 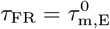 given in Section 4.3.
- Connections: *I*_0_ = −20pA, *I*_1_ = 43pA, *J*_0_ = −0.35pA/Hz, *J*_1_ = 2.7pA/Hz were calculated by Eq 44.
- Numerical parameters: ring was discretized by 40 points; time step 0.02ms.

## 5 Acknowledgements

Erez Persi contributed valuable insights during the development of this work, and the authors also thank Nicolas Gazères for reviewing an earlier version of the manuscript.

